# An M-learner approach for heterogeneous mediation analysis with high-dimensional omics mediators

**DOI:** 10.64898/2026.08.25.747106

**Authors:** Xingyu Li, Peng Wei

## Abstract

Causal mediation analysis is widely used to identify biological pathways linking exposures to outcomes, but most methods assume homogeneous mediation effects across individuals. In high-dimensional omics settings, this assumption can mask important heterogeneity driven by demographic, genetic, or environmental factors. We propose the **M-high-learner**, a flexible framework for detecting heterogeneous mediation effects with high-dimensional mediators. The method identifies mediators with subgroup-specific indirect effects while distinguishing them from null or homogeneous signals and controlling the type I error rate. It is computationally efficient, scalable, and yields interpretable sub-types. Simulation studies show that the proposed approach achieves high power while maintaining accurate error control. Applications to the Framingham Heart Study and the Multi-Ethnic Study of Atherosclerosis reveal that the mediation role of gene expression in sex’s effect on high-density lipoprotein varies across subgroups defined by body mass index and age. Our framework provides a practical tool for uncovering heterogeneous biological mechanisms in high-dimensional genomic studies.

**Author Summary:** Biological processes linking risk factors to disease often differ across individuals, but many existing methods assume these processes are the same for everyone. This can hide important differences between groups. We developed a powerful method to identify when these pathways vary across subgroups using large-scale molecular data. Our approach detects differences in how intermediate biological factors contribute to outcomes in populations defined by characteristics such as age and body mass index. Applying our method to population studies, we found that some biological pathways operate differently across groups, suggesting that key mechanisms may be missed when differences are ignored. Our work provides a tool to better understand how disease-related processes vary across individuals, which may support more targeted and personalized approaches to health research.

## Introduction

Causal mediation analysis is a widely used framework in genomics, as it enables researchers to investigate how exposures influence traits through intermediate biological pathways, thereby providing mechanistic insight into exposure-intermediate molecular traits-phenotype relationships [1, 2, 3]. However, most existing studies assume homogeneous mediation effects of intermediate omics traits, e.g., gene expression, on the pathway from an exposure to a phenotype across individuals, effectively treating the population as a single, uniform group [4, 5].

In practice, mediation effects are often heterogeneous due to variation in demographic and genetic characteristics, medical history, lifestyle factors, and other unobserved attributes. Mediators may differ across subpopulations or exert distinct effects on the outcome under different biological or environmental conditions. For example, high-density lipoprotein (HDL) levels are associated with both mass index (BMI) and sex, and the relationship between BMI and HDL has been shown to vary by sex [6]. This phenomenon has been consistently documented in epidemiological and clinical studies, where elevated BMI is negatively associated with HDL levels, potentially through mechanisms involving insulin resistance, increased triglycerides, and inflammation in adipose tissue. In such settings, assuming homogeneous mediation effects may obscure biologically meaningful pathways, as opposing effects across subtypes can cancel each other out and mask true mediation signals.

In high-dimensional genomic settings, the effects of molecular mediators are unlikely to be uniform across the population, as regulatory pathways may contribute differently to disease processes under varying genetic and environmental contexts. Accounting for heterogeneity in mediation effects is therefore essential for accurately characterizing the functional roles of candidate mediators and for identifying subpopulations in which specific molecular mechanisms drive phenotypic variation. Ignoring such heterogeneity may lead to incomplete or misleading interpretations of genomic data, particularly when signals are present only in a subset of individuals.

In recent years, several statistical methods have been proposed to address heterogeneous mediation effects [7, 8, 9]. However, these approaches rely on predefined subgroups or are limited to settings involving a single mediator. In contrast, the method proposed by [10] determines subgroup membership through a penalty term in the loss function, but requires the number of subgroups to be specified in advance and does not provide a formal mechanism to test for the presence of heterogeneity.

[11] introduced the M-learner framework to address heterogeneous mediation analysis. Nevertheless, the M-learner is limited to single-mediator settings and is not readily applicable to settings involving multiple or high-dimensional mediators. To overcome these limitations, we propose the M-high-learner, a novel framework designed to accommodate multiple mediators and operate effectively in high-dimensional settings.

Our method addresses heterogeneity in high-dimensional mediation analysis by identifying mediators that exhibit subgroup-specific indirect effects while excluding both non-heterogeneous and globally heterogeneous cases. The proposed approach controls the type I error rate and yields interpretable subgroups that reflect underlying biological heterogeneity. It is computationally efficient, flexible, and scalable, while maintaining statistical power in complex settings. Through extensive simulations, we demonstrate that the method reliably detects heterogeneous subgroups driven by mediator-specific indirect effects while correctly rejecting non-heterogeneous scenarios. Applications to the Framingham Heart Study (FHS) and the Multi-Ethnic Study of Atherosclerosis (MESA) further illustrate that sex influences HDL through heterogeneous pathways of gene expression, with BMI and age emerging as key determinants of this variability across populations.

## Description of the Methods

In this section, we introduce the mediation models and relevant notations, then we introduce the proposed framework, M-high-learner.

### Mediation Analysis

Mediation analysis is a statistical approach used to investigate the mechanism through which an independent variable (*W* ) affects a dependent variable (*Y* ) via a series of mediator (*M* ). In mediation analysis, researchers are not only interested in the total effect of *W* on *Y* but also in the indirect effect transmitted through specific mediators and the direct effect that bypasses the mediators. A mediator is a variable that conveys part of the influence of the independent variable on the dependent variable, thereby revealing the underlying mechanism of action. In the genetic and genomic context, we focus on how exposures influence biological traits through potentially high-dimensional omics mediators, e.g., gene expression, DNA methylation or protein expression [6, 12]. In this study, the exposure of interest is sex whereas the outcome of interest is HDL. Given the complexity of biological relationships, there are often multiple mediators, namely different genes’ expression levels, denoted as *M* ^(1)^, …, *M* ^(*k*)^. In the absence of heterogeneity assumptions, the mediation effects are typically evaluated using linear models.

A mediation model (Figure 1) consists of the following equations. Without loss of generality, we assume the outcome, and mediator variables are standardized to have mean 0 and variance 1, the exposure has two levels 0 and 1.

**Figure 1.**
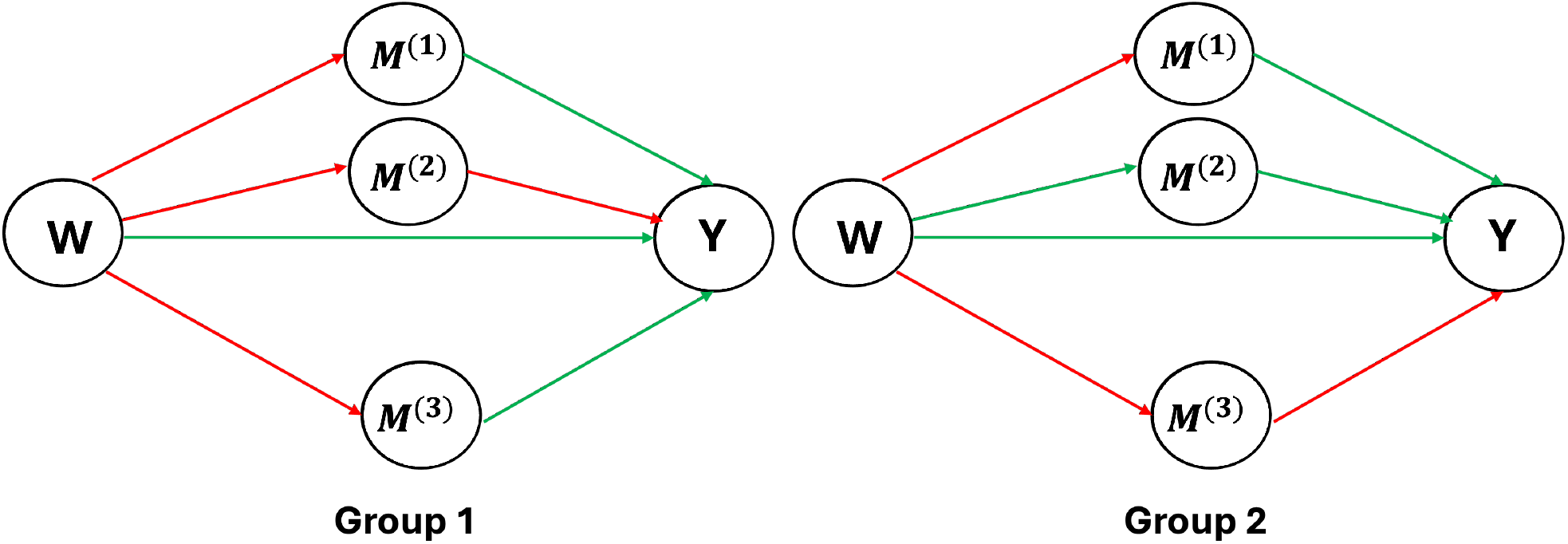
Directed Acyclic Graph. The figure illustrates the relationship among W, M, and Y within different heterogeneous groups. The effect from W to Y through M is the indirect effect, while the effect directly from W to Y is the direct effect. The red color indicates a positive effect, and the green color indicates a negative effect. The two groups represent different heterogeneous subgroups.

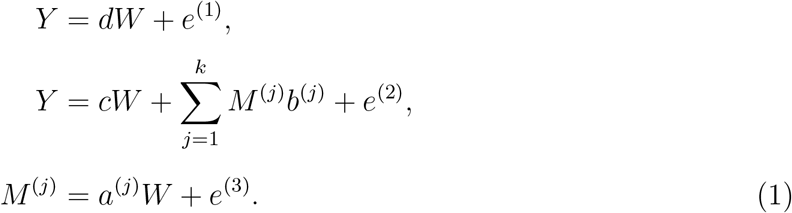

In (1), *k* is the total number of mediators. When *k* = 1, it corresponds to a single mediator model; otherwise, it corresponds to a multiple-mediator model. *Y* is the continuous dependent (outcome) variable; *W* is the independent (exposure) variable; *M* ^(*j*)^ is the *j*th mediator; *e*^(1)^, *e*^(2)^, *e*^(3)^ are residuals for each equation; *a*^(*j*)^, *b*^(*j*)^, *r* and *c* are regression coefficients, typically estimated by the maximum likelihood estimation (MLE) method. The parameter *d* is the total effect and *c* is the direct effect.

A common approach to quantify the mediation effect is the product measure [13], which estimates the indirect effect as the product of the effect of *W* on the mediator (typically denoted as *a*) and the effect of the mediator on *Y* (typically denoted as *b*), i.e. product 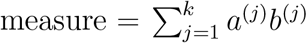 . This method is intuitive, easy to interpret, and widely used in structural equation modeling (SEM) and regression frameworks. By applying the product measure, mediation analysis allows a quantitative assessment of the extent to which an independent variable influences a dependent variable through mediators, providing a more comprehensive understanding of the causal relationships among variables.

Recently, another measure is proposed [14, 15, 16], called 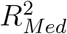, which is an appealing complementary measure to traditional total mediation effect measures, such as the product measure, by avoiding the issue of cancellation from component-wise mediation effects *a*^(*j*)^*b*^(*j*)^’s of different directions [17, 18].

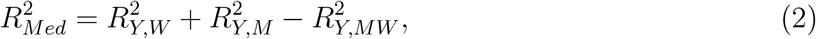

where 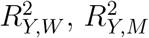 and 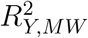 represent the coefficient of determination for the regression 2 models in which *Y* is regression on *W, M* and (*W, M* ), respectively. In addition, there *Med* is another measure Shared Over Simple (SOS), which is defined as 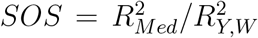. SOS is a relative measure of 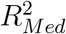; it is the standardized exposure-related variance in the outcome that is shared with the mediator.

Under model (1), all three measures, product measure, 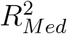, and SOS, ignore heterogeneity; that is, the effect of the exposure on the outcome through the mediator is assumed to be independent of covariates, implying that the mediated effect represents an average effect across all individuals. These methods evaluate mediation effects at the global level without accounting for heterogeneity, and therefore cannot identify mediators with heterogeneous mediation effects. To this end, we introduce a novel, flexible, powerful, computationally efficient, and interpretable machine learning–based approach to uncover the complex relationships among covariates *X*, mediators *M*, exposure *W*, and outcome *Y* in high-dimensional mediation analysis.

### Setup and Assumptions

Let 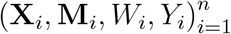 be independent and identically distributed samples from the distribution of (**X, M**, *W, Y* ), where *X* = (*X*^(1)^, …, *X*^(*p*)^) is a p-dimensional vector of covariates, **M** = (*M* ^(1)^, …, *M* ^(*k*)^) is the intermediate variable (mediators), *k* is the dimension of mediators, *W* = *w* ∈ {0, 1} be the binary exposure variable.

We adopt the potential outcomes framework under the Stable Unit Treatment Value Assumption (SUTVA) [19], and let **M**(*w*) be the potential values of the intermediate variable if the unit were to receive exposure condition *w. Y* (*w*, **M**(*w*^*∗*^)) denotes the counterfactual outcome that would have been observed if the exposure *W* were set to *w* and the mediator **M** to the value **M**(*w*^*∗*^). In addition, we assume the sequential ignorability, meaning that, conditional on the observed covariates, there is no unmeasured confounding of the exposure–mediator and mediator–outcome relationships [20].

For outcome *Y*, mediator **M** and covariates **X**, we assume

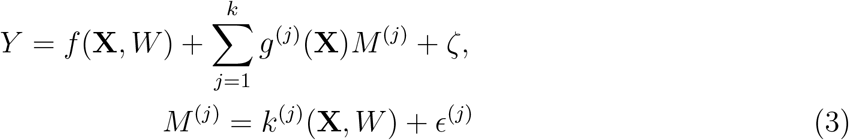

where *f, g*^(*j*)^, *k*^(*j*)^ can be arbitrary integrable functions, *ζ* and *ϵ*^*j*^ are the error terms with zero mean. In (3), function *f* represents the direct effect, 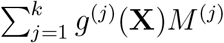 represents the indirect effect.

This condition is not very restrictive; for example, the following linear structures (4) satisfies the expression (3):

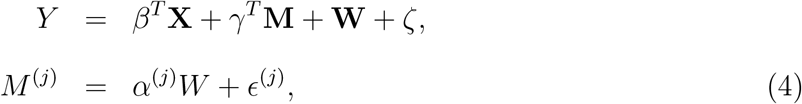

where *β* ∈ ℝ^*p*^ and *γ* ∈ ℝ ^*k*^ are the coefficients vectors in (4).

In traditional mediation analysis, the average total effect (ATE), average indirect effect (AIE) and average direct effect (ADE) are of interest, which are defined as

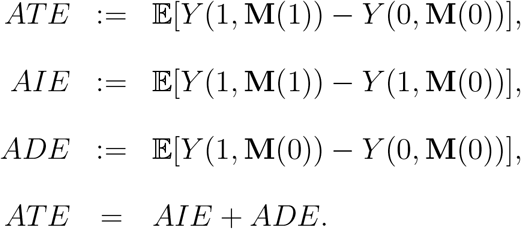

However, these effects reflect the average effect across all individuals and cannot capture heterogeneity. In order to explore mediation heterogeneity, we focus on the indirect effect conditional on covariates, which is denoted as conditional average indirect effect (CAIE), defined as

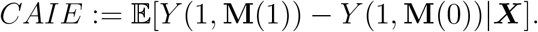

CAIE quantifies the indirect effect (IE) conditional on covariate ***X*** and thus it can capture the heterogeneity.

Subgroups by heterogeneous IEs can be defined as

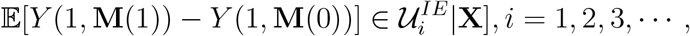

where 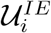 is the set, for any 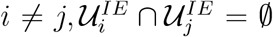, the number of sets are unknown, and when there is no heterogeneity, the number of the set is 1.

### Proposed method: M-high-learner

We propose a novel algorithm, termed M-high-learner, to address the estimation of heterogeneous mediation effects in high-dimensional settings. The framework consists of three main steps (see Figure 2 for an illustration of the pipeline and Appendix B for the detailed pipeline of M-learner):

**Figure 2.**
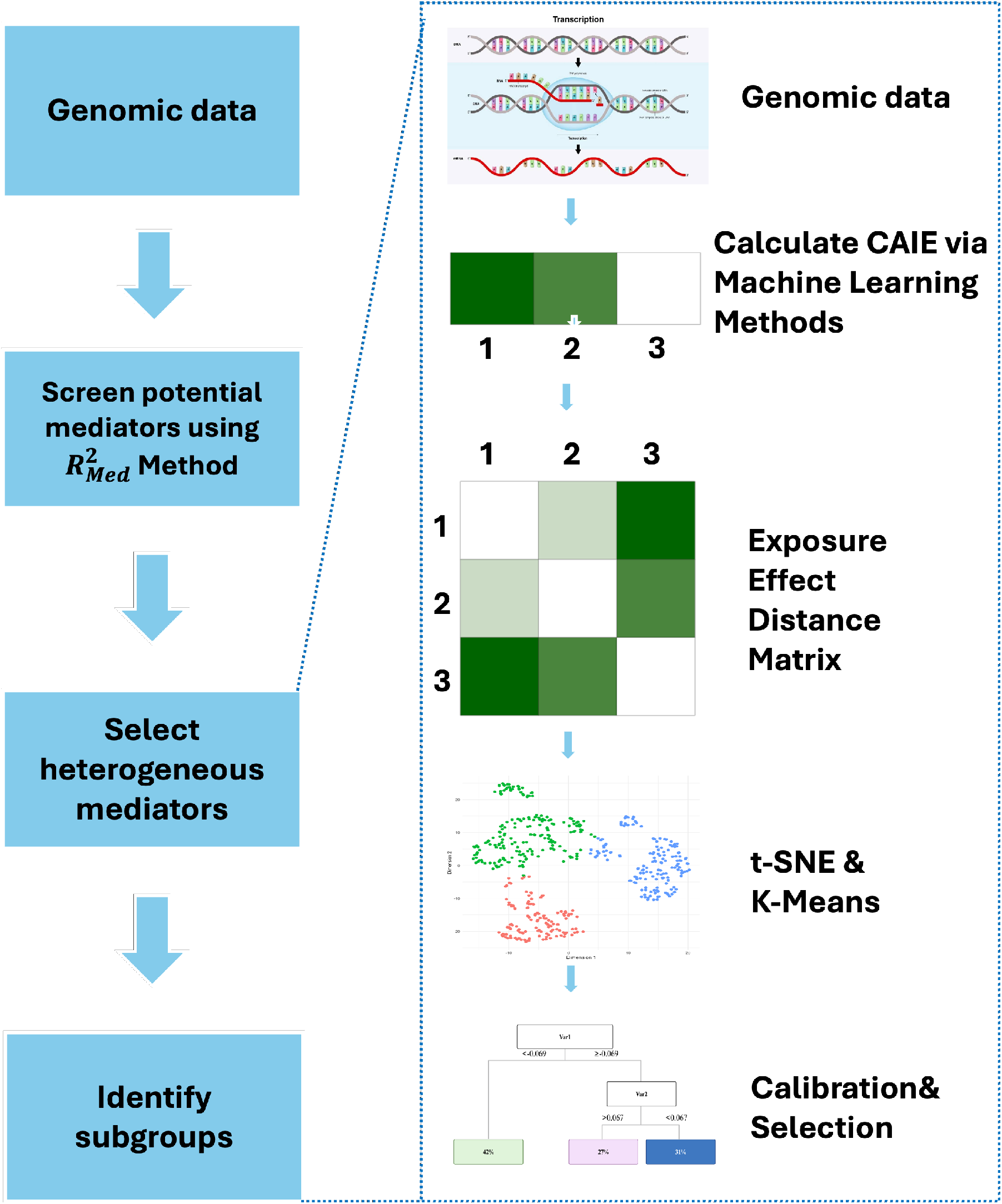
Overview of the M-high-learner pipeline for identifying heterogeneous mediation effects from high-dimensional genomic data. Potential mediators are first screened using the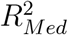 method. Heterogeneous mediators are then selected using the M-learner in a single-mediator setting. The selected mediators are subsequently used to estimate conditional average indirect effects (CAIE) and derive a exposure effect distance matrix. Finally, subgroup identification is performed via modified t-SNE and K-means clustering, and the resulting subgroups are calibrated and interpreted using a decision tree.

1. Filter potential mediators with the 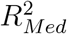 -based mediator selection procedure.
2. To further screen mediators with heterogeneous effects, M-learner method is applied to each mediator individually.
3. M-learner is applied to filtered mediators to identify heterogeneous subgroups.

Here, we introduce the proposed method in detail.

Step 1. The first step is to screen potential mediators. In genomic studies, the number of candidate mediators can be on the order of tens of thousands, whereas the sample size is usually only a few thousand. Therefore, an initial screening step to reduce the number of mediators is necessary. We adopt the method proposed in [14] to select potential mediators.

Step 2. For the mediators retained from the first step, we apply the single-mediator M-learner approach to examine whether each mediator exhibits heterogeneous effects. Only those mediators with evidence of heterogeneity are kept for further analysis.

Step 3. Finally, for the mediators retained in Step 2, we apply the multi-mediator M-learner approach to detect heterogeneous subgroups.

### Estimating CAIE via M-high-learner

Specifically, we use M-high-learner to estimate the CAIE:

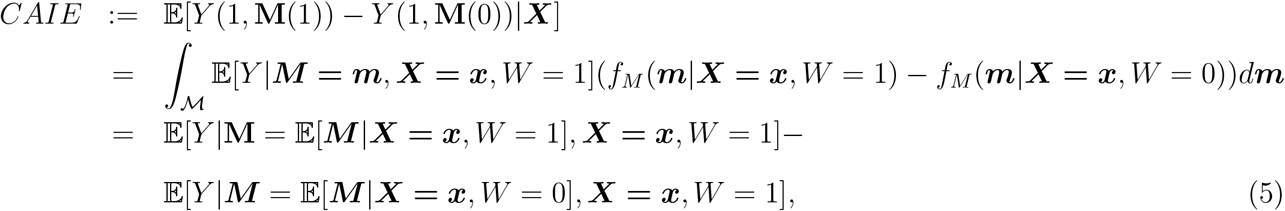

where *M* denoting the space of the mediator ***M***, *f*_*M*_ (·) is the conditional density function of ***M*** .

The estimation consists of three steps.

First, the mediator function is estimated conditional on the exposure being at level 1,

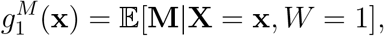

with a base learner, using the observations {(**M**_*i*_, *Y*_*i*_)}_*Wi*_=1 and denoting the estimator 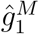 [21]. This step is to estimate the mediators for the exposure at level 1. The base learner can be any machine learning models. However, when there are multiple mediators, we recommend using random forests, as they can account for the correlations among mediators.

Second, the mediator function is estimated conditional on the exposure being at level 0,

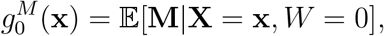

with a base learner, using the observations {(**X**_*i*_, *Y*_*i*_)}_*Wi*_=0 and denoting the estimator by 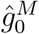. This step is to estimate the mediators for the exposure at level 0.

Finally, fit the response function on the exposure being at level 1,

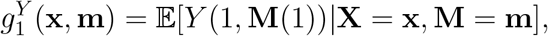

with a base learner, using the observations {(**X**_*i*_, **M**_*i*_, *Y*_*i*_)}_*Wi*_=1, and denoting the estimator by 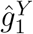 . This step is used to fit the relationship among the covariates **X**, mediators **M** and outcome *Y* .

### Subgroup Identification via Similarity-Based Clustering

The M-high-learner then leads to

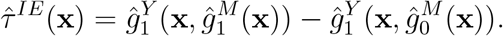

Now that we have estimated the CAIE, it remains to identify subgroups of subjects. We propose a new method to transform the estimation of CAIE to the clustering. The estimated CAIE for each subject *i* is denoted by 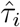.

First, we evaluate the CAIE difference between each pair of subjects *i* and *j*, which we refer to as the exposure effect distance. It is defined by *dis*(*i, j*), the distance metric can be defined as Euclidean distance, Manhattan distance, or other formulations. This exposure effect distance reflects the difference in exposure effects between two subjects. A smaller distance between the exposure effects of two subjects indicates greater similarity between them. Conversely, when the indirect effects differ substantially, the distance will be larger, reflecting lower similarity between the two subjects. Thus, the exposure effect distance serves as a measure of similarity in exposure effects; see Figure 2 for an illustration. In the following analysis, we consistently adopt the Euclidean distance to compute the distance between subjects *i* and *j* which means 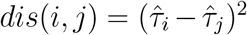. Consequently, an *n* × *n* exposure effect distance matrix is obtained. This matrix is then projected into a two-dimensional Euclidean space using t-SNE [22], where each subject *i* is assigned a coordinate, the detailed steps of t-SNE can be found in the Appendix A (in proposed method, the t-SNE is different from the original t-SNE method in [22], we introduce the modified t-SNE method in the Appendix). The objective of this step is to preserve the local similarity of the indirect effect distance between subjects *i* and *j*. As demonstrated in [23], under appropriate conditions, t-SNE possesses desirable theoretical properties for effectively maintaining local similarity. In addition, [24, 25] provide further discussion. Empirically, t-SNE has been shown to be one of the most effective methods for preserving local neighborhood relationships.

Subsequently, K-means clustering is performed on the projected points in the Euclidean space. The range of cluster numbers (from 2 to *K*) for K-means clustering must be specified in advance, based on the sample size and the number of covariates; an excessively large *K* can result in substantial overfitting. Based on our experience, In this study, we set 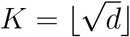, where *d* denotes the number of covariates and ⌊·⌋ denotes the floor function. A detailed discussion of parameter *K* and reason for K-means can be found in the Discussion section.

Then, a decision tree is employed to model the clustering results obtained for each predefined number of clusters, with the objective of mapping the unknown categories to interpretable categorical information. For each leaf of the decision tree, we refer to it as a subgroup. To mitigate overfitting, we impose three constraints on decision-tree construction: (i) a minimum leaf size, which specifies the minimum number of individuals required in each subgroup; (ii) a maximum tree depth; and (iii) a restriction that prevents the same variable from being used in multiple splits within a tree. Further details and methodological justification are provided in Appendix B. We use *p*_*leaf*_ to select the final, unique subgroups from different decision tree results. Each 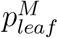 is derived from multiple test statistics. Specifically, we compute 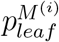 for each mediator, and subsequently aggregate them using the Aggregated Cauchy Association Test (ACAT) to obtain the final 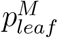 [26] with multiple mediators. When there is only one mediator, we use 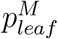 directly.

The leaf-level heterogeneity measure 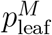 is defined as follows. For the *i*th mediator, we compare the following two nested models:

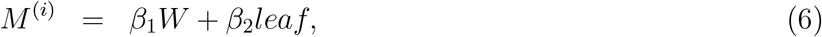

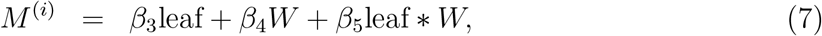

Let *L*_0_ and *L*_1_ denote the likelihoods corresponding to models (6) and (7), respectively, and let *G* denote the number of leaves in the decision tree. Under the null hypothesis of no exposure-effect heterogeneity across leaves, the likelihood-ratio statistic (LRT), 2(log *L*_1_ − log *L*_0_) asymptotically follows a chi-squared distribution with (G-1) degrees of freedom.

The corresponding p-value is denoted by 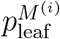. Among the candidate mediators, we define 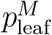 based on the mediator yielding the strongest evidence of heterogeneous exposure effects. Similarly, replacing *M* with the continuous outcome *Y* gives 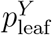.

Compared with the definition of *p*_leaf_ proposed by [25], we modify the formulation to improve its robustness as the number of candidate covariates increases. With increasing covariate dimensionality, the number of potential data-adaptive partitions also increases substantially. Consequently, the selected leaves may exhibit differences in their mean mediator or outcome levels simply as a result of the enlarged search space, even in the absence of true exposure-effect heterogeneity. If the leaf main effect is omitted from the null model, the likelihood-ratio test may therefore capture both differences in the mean response across leaves and differences in the exposure effect across leaves. To better isolate exposure-effect heterogeneity, we include the leaf main effect in both the null and alternative models. Under this specification, the likelihood-ratio test specifically evaluates the additional contribution of the exposure-by-leaf interaction after accounting for the main effect of leaf membership. Our simulation studies further support this modification, demonstrating improved calibration as the number of candidate covariates increases. The calibrated thresholds 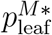 and 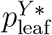 are used to screen candidate profiles for evidence of heterogeneous exposure effects on both the mediator and the outcome. Among the candidate leaf profiles satisfying both calibration criteria, the profile with the smallest 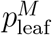 the final subgroup classification. Specifically, the selected profile is defined as is selected as

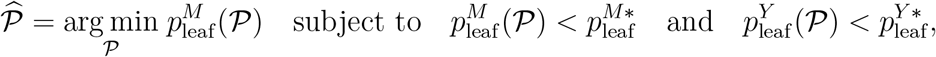

where *P* denotes a candidate profile, where 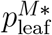 and 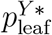 are determined through the calibration procedure. If no candidate profile satisfies both calibrated criteria, no heterogeneous subgroup structure is identified.

The calibration step plays a critical role in controlling spurious subgroup identification arising from the enlarged, data-adaptive search space. In particular, as the number of candidate covariates increases, the algorithm has more opportunities to identify apparently heterogeneous partitions by chance, even under the Global and Null scenarios where no true exposure-effect heterogeneity is present. The calibrated thresholds therefore provide an empirical safeguard against such false discoveries, enabling the procedure to distinguish genuine heterogeneous subgroup structures from chance patterns induced by extensive subgroup searching.

In our method, we employ base learners to estimate various functions. The base learners can be random forests, XGBoost, neural networks, BART, among others [27, 28, 29, 30]. However, when dealing with high-dimensional mediators, we recommend using random forests as the base learner. This is because random forests account for the correlations among multivariate responses when making predictions, whereas many other learners, such as XGBoost, typically predict each response variable independently and thus fail to capture their mutual dependencies [30].

### Simulations

To evaluate the performance of the proposed M-high-learner framework, we design different scenarios that reflect varying real-world heterogeneity structures. Our simulation comprises two components: one involving a single mediator and the other involving multiple mediators. We describe each part in the following sections.

### Single-mediator settings

In experiment 1-5, the sample size was fixed at 1, 000, with 50 covariates generated for each subject *i*. In Experiments 6 through 10, we systematically increased the sample size from 1, 000 to 2, 000 to evaluate how the performance of the proposed method varies with increasing data availability.

Specifically, we simulated 10 scenarios; (1) existing heterogeneity, all effects via the mediator (All); (2) existing heterogeneity, part of the effect via mediator (Part); (3) no heterogeneity, all units benefit from the exposure (Global); (4) no heterogeneity, 0% effects via mediator (Null 1); (5) no heterogeneity, 0% effects via mediator (Null 2). The detailed set up can be found in Appendix C.

We calibrated the thresholds based on Scenario Null1 and Null2 by controlling the Type I error at 10%, and then applied this thresholds to assess the validity of subgroup identification across other scenarios.

From the Figure 3, Figure 5, Figure 6, and Appendix Figure 4, we can observe that M-high-learner can effectively and accurately identify subgroups exhibiting heterogeneity in the indirect effect, regardless of whether one or multiple mediators are involved. Moreover, it adequately controls the type I error rate when either the entire effect or only a partial effect passes through the mediators. From the distribution of the identification thresholds in Figure 5, the results appear to be conservative.

**Figure 3.**
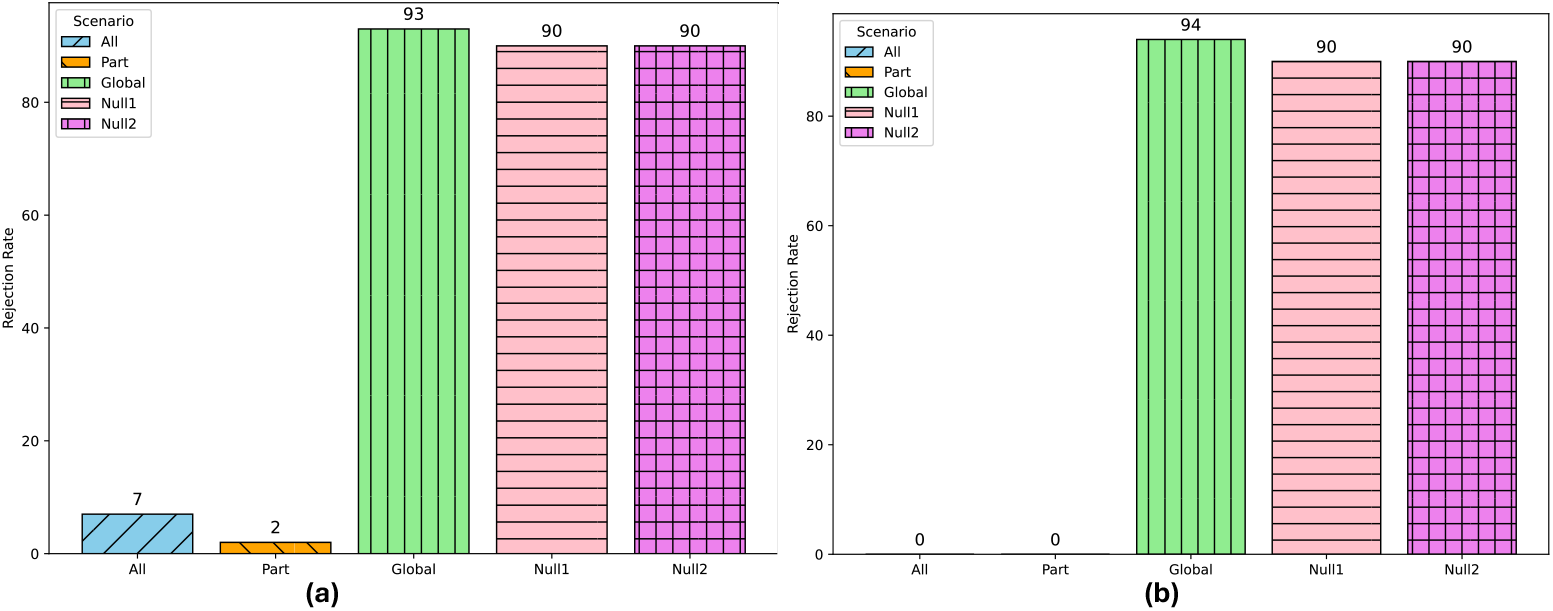
Calibrated rejection rate for single-mediator settings. Calibrated rejection rate: proportion of 100 simulations in which the null hypothesis of no heterogeneous indirect effect was rejected after calibration, reflecting the method’s power to detect mediation heterogeneity while controlling type I error. (a) Sample size was 1000, (b) Sample size was 2000.

**Figure 4.**
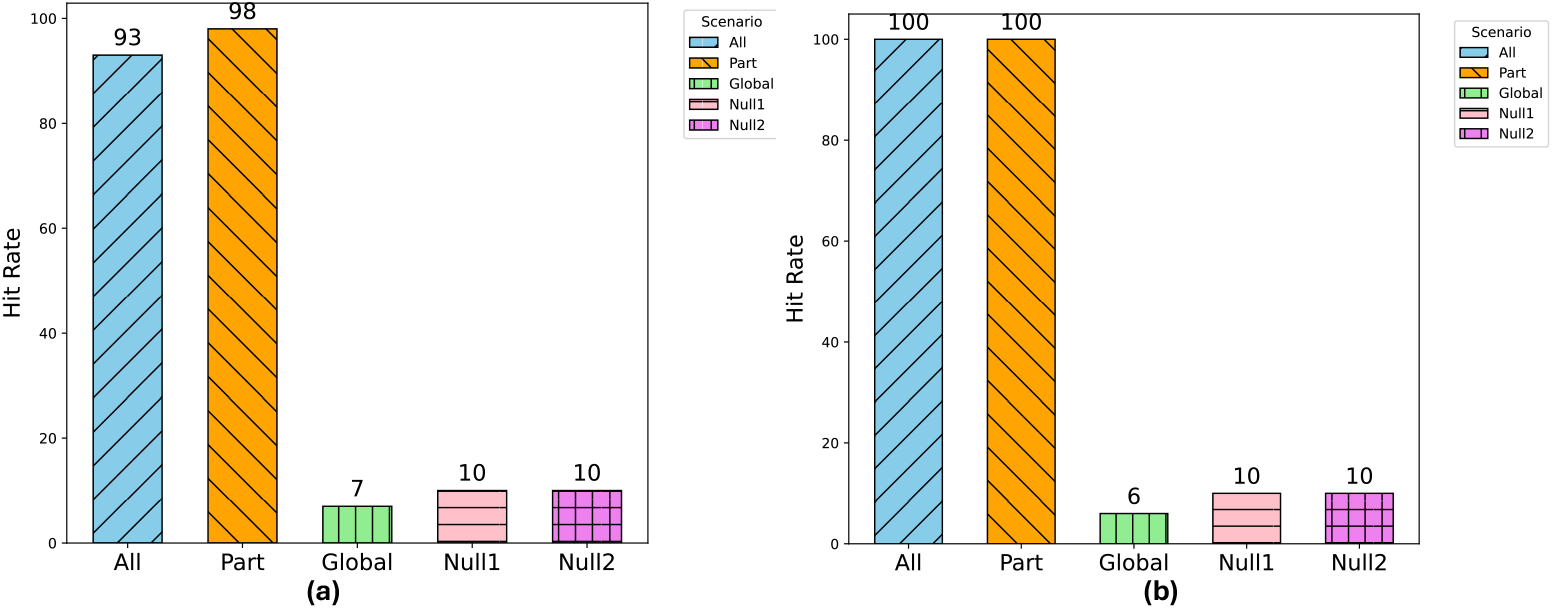
Proportion of 100 simulation runs in which the method correctly identified the heterogeneous variables *X*^(1)^ and *X*^(2)^. (a) Sample size was 1000, (b) Sample size was 2000.

**Figure 5.**
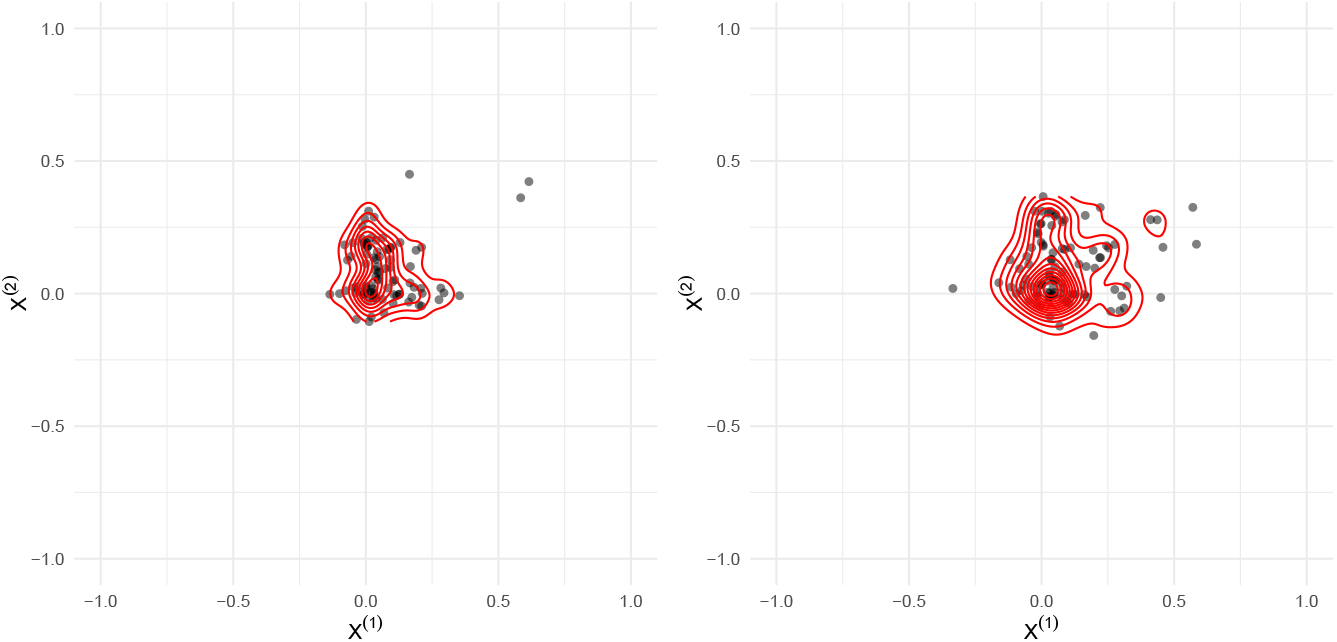
Threshold distributions for single-mediator settings when sample size was 1000. Solid dots represent the estimated thresholds of heterogeneous subgroups obtained in each experiment, with values greater than 1 or undefined thresholds set to 1. The red line depicts the density curve. The left panel is All scenario, the right panel is Part scenario.

**Figure 6.**
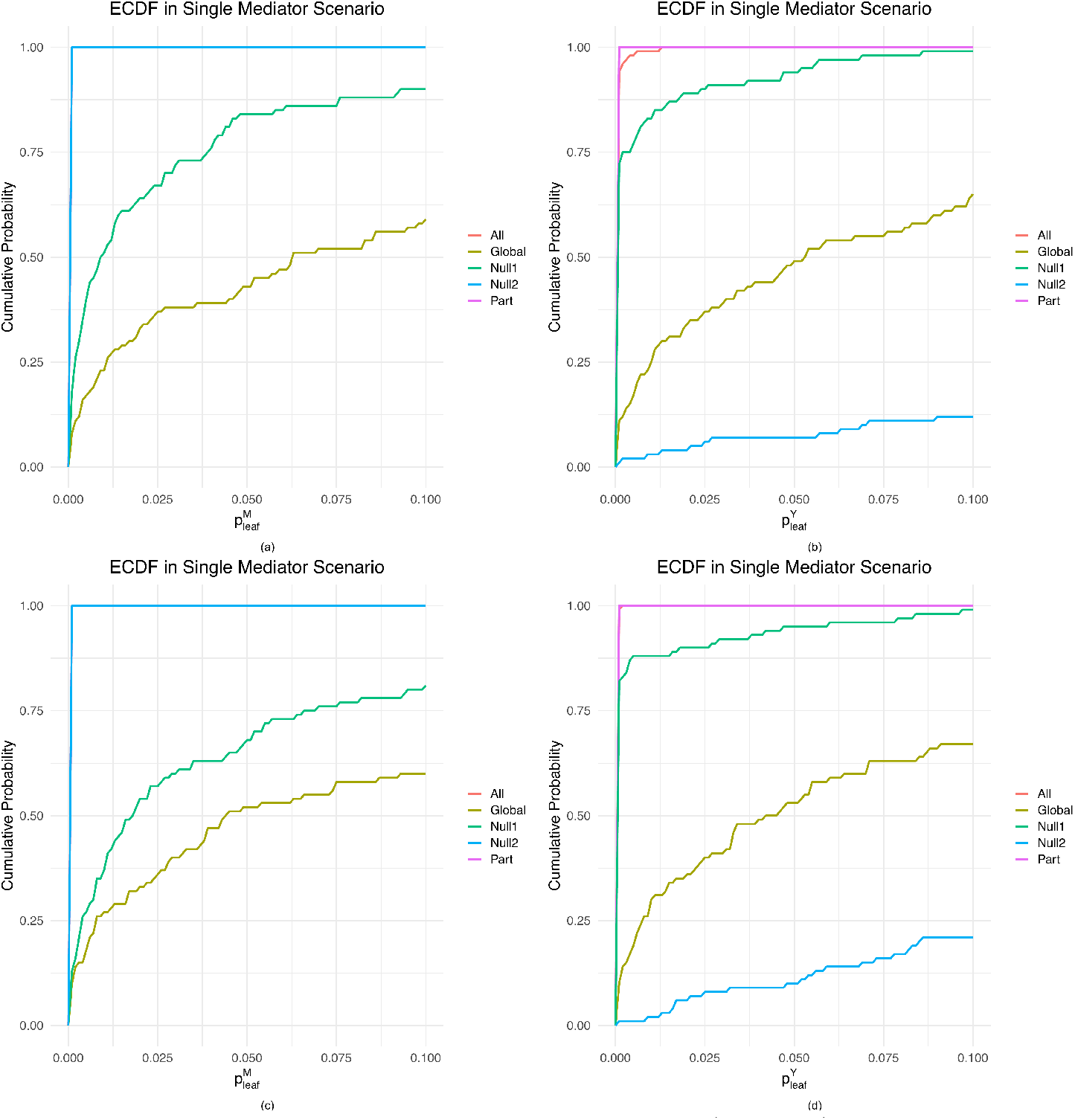
Empirical cumulative distribution functions (ECDF) of *p*_*leaf*_ under single mediator scenario. (a) 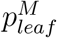 when sample size was 1000, (b) 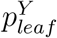 when sample size was 1000, (c) 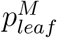 when sample size was 2000, (d) 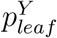 when sample size was 2000.

### Multiple-mediator settings

In experiment 10 − 15, the sample size was fixed at 1, 000, with 50 covariates generated for each subject, 10 mediators, and all 10 mediators were associated with exposure . In Experiments 16 through 20, we systematically increased the sample size from 1, 000 to 2, 000 to evaluated how the performance of the proposed method varies with increasing data availability.

Specifically, we simulated 10 scenarios, (1) heterogeneity present, all effects via the mediator (All); (2) heterogeneity present, part of the effect via mediator (Part); (3) no heterogeneity, the exposure has equal mediation effects on all subjects (Global); (4) no heterogeneity, 0% effects via mediator (Null 1); (5) no heterogeneity, 0% effects via mediator (Null 2). The detailed set up can be found in Appendix C.

Similar to the single-mediator settings, we used Null1 and Null2 for calibration; however, we adopted a more conservative strategy to control the type I error rate at 5%.

From Figure 7 and Figure 8, Appendix Figure 5 and 6, it can be observed that the M-high-learner effectively and accurately identifies subgroups exhibiting heterogeneity in the indirect effect across multiple mediators. These results demonstrate that the proposed method can reliably detect heterogeneous regions even in high-dimensional settings involving multiple mediators.

**Figure 7.**
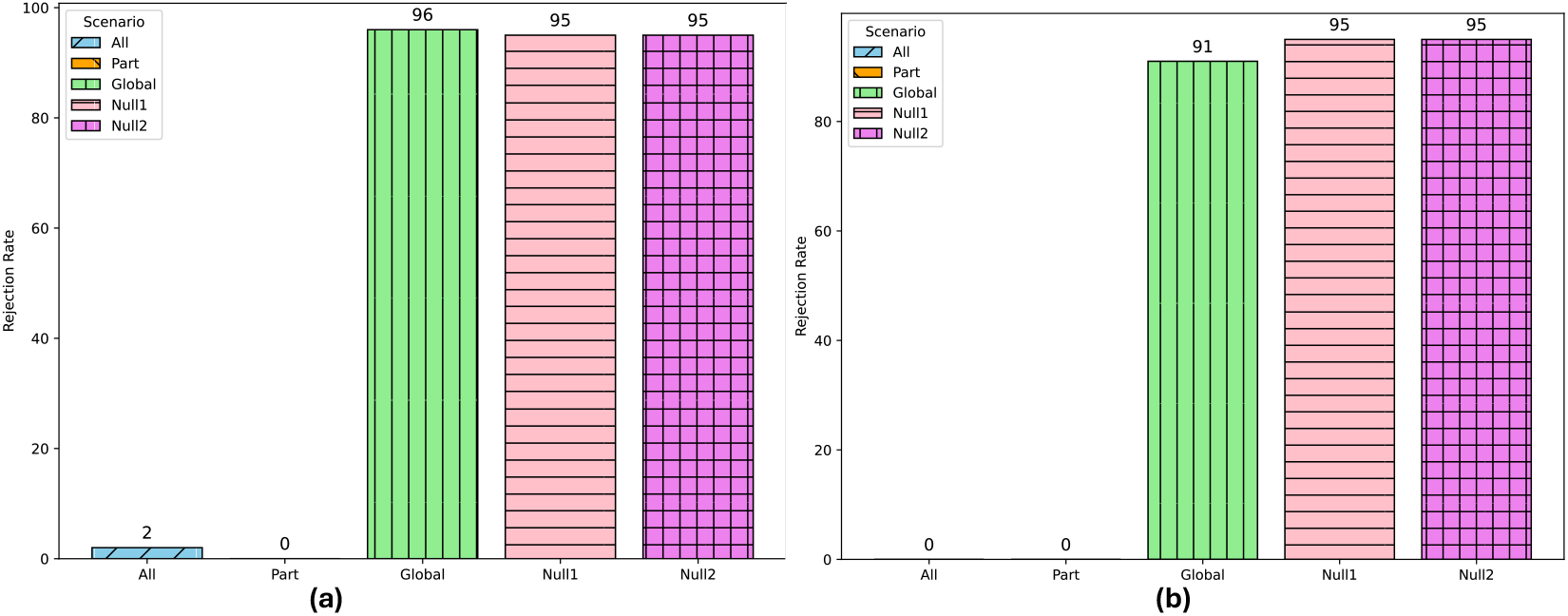
Calibrated rejection rate for multiple-mediator settings. Calibrated rejection rate: proportion of 100 simulations in which the null hypothesis of no heterogeneous indirect effect was rejected after calibration, reflecting the method’s power to detect mediation heterogeneity while controlling type I error. (a) sample size was 1000, (b) sample size was 2000.

**Figure 8.**
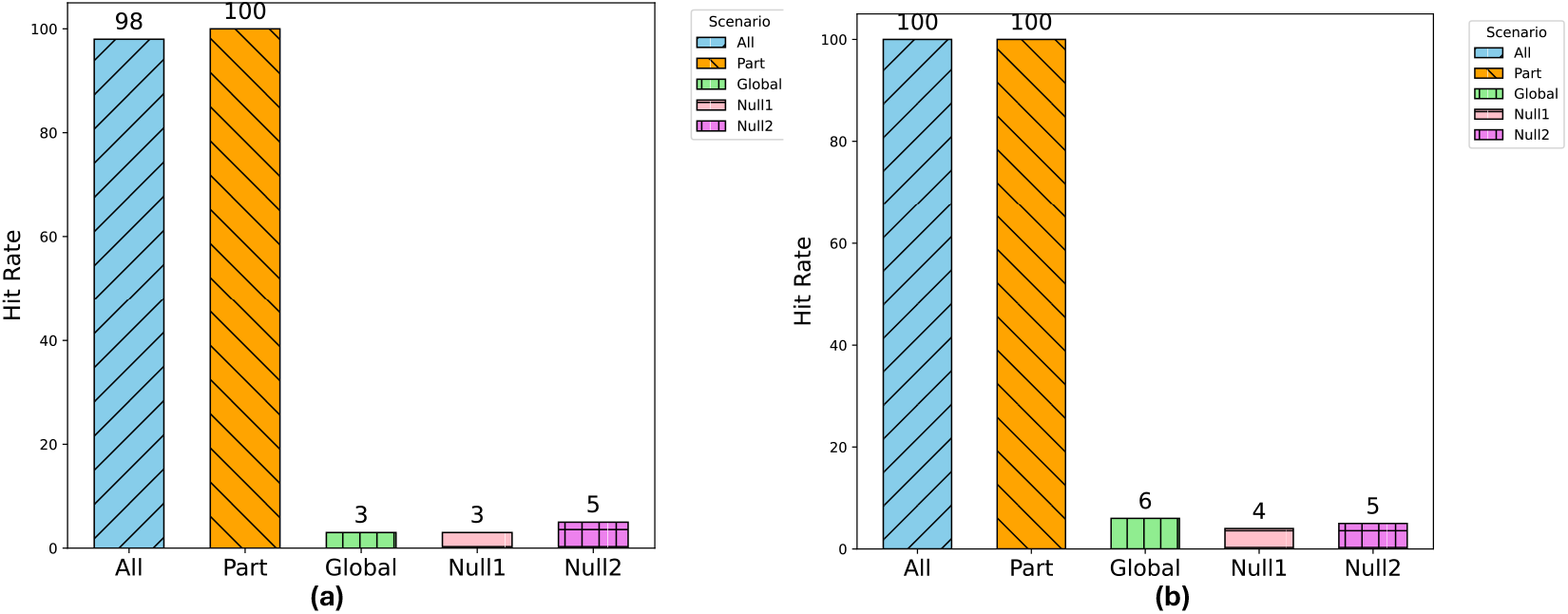
Proportion of 100 simulation runs in which the method correctly identified the heterogeneous variables *X*^(1)^ and *X*^(2)^ for multiple mediators. (a) Sample size was 1000, (b) Sample size was 2000.

The simulation results for both single-mediator and multiple-mediator settings demonstrate that the proposed method can reliably identify heterogeneous mediators.

## Application to the FHS and MESA studies

To demonstrate the performance of the proposed M-high-learner method in real-world datasets, we analyzed the Framingham Heart Study (FHS) to identify subgroups showing heterogeneous mediation effects of gene expression on the pathway from sex to HDL, and further replicated the findings in the Multi-Ethnic Study of Atherosclerosis (MESA) study. The Trans-Omics for Precision Medicine (TOPMed) program, initiated by the National Heart, Lung, and Blood Institute (NHLBI), is a large-scale effort to integrate whole-genome sequencing (WGS) and other omics data across more than 85 population-based studies [31, 32]. The program aims to elucidate the genetic and molecular mechanisms underlying disorders of the heart, lung, blood, and sleep [32].

Within TOPMed, the FHS recruited the Offspring cohort in 1971, comprising the children of the Original cohort and their spouses. This cohort includes 5, 124 individuals, 52% of whom are female [33]. In 2002, the FHS launched the Third-Generation cohort, consisting of 4, 095 participants, 54% female, who are the children of the Offspring cohort [34]. Another key cohort within TOPMed is the MESA study, which enrolled 6, 814 participants aged 45–84 years old from six U.S. communities [35]. MESA focuses on identifying risk factors for cardiovascular disease (CVD), particularly atherosclerosis, across four ethnic groups: Non-Hispanic Whites, African Americans, Hispanics, and Chinese Americans [36]. The transcriptome encompasses all messenger RNAs (mRNAs) or transcripts present in a cell under a specific condition or developmental stage. Transcriptomic profiling is critical for interpreting the functional elements of the genome, characterizing cellular components, and understanding development and disease [34]. While hybridization-based microarray gene expression profiling remains cost-effective and high-throughput, it is limited by current genomic knowledge [37]. In contrast, RNA sequencing (RNA-seq) enables the discovery of novel gene transcripts and non-coding RNAs [38]. With the declining cost of next-generation sequencing technologies, RNA-seq has become increasingly feasible for large-scale studies, such as FHS and MESA.

Previous research have highlighted notable sexual dimorphism in HDL cholesterol levels and functionality [39, 40]. We applied our developed M-high-learner method to the FHS study as the discovery cohort (n=4472 with microarray-based gene expression profiling) and the MESA study as the replication cohort (n=1124; RNA seq-based gene expression profiling) to estimate heterogeneous indirect effects due to gene expression and identify heterogeneous subgroups underlying sex-related variation in HDL. HDL was measured from EDTA plasma (mg/dL), and age was recorded at the time of examination.

Covariates included body mass index (BMI, kg/m^2^), dichotomized smoking status (current smoker vs. non-smoker), dichotomized drinking status (never vs. ever), subcohort indicator for the FHS (Offspring or Third Generation), and ethnicity for the MESA. Additionally, the top 10 principal components (PCs) of genome-wide gene expression data, selected based on eigenvalues, were included to account for population structure in the mediation analysis models. The use of PCs is standard in genome-wide association studies, as they help correct for subtle population stratification and control for confounding genetic backgrounds [6, 15, 41].

As shown in Figure 9 (a), M-high-learner identified four subgroups in the FHS. The M-high-learning model selected BMI and age as the baseline covariates to define subgroups in the FHS. Specifically, it selected 25 as the threshold for BMI, which is consistent with the clinical definition of overweight [42, 43]. In addition, the model selected 50 and 62 years old as the thesholds for age. Among individuals with lower BMI, women typically experience menopause around the age of 50, during which the decline in estrogen substantially alters HDL metabolism, including particle remodeling and reduced antioxidant capacity [44, 45]. In contrast, among individuals with higher BMI, after the age of 62, sex-related hormonal differences exert a stronger influence on HDL secretion and regulation. Overweight and Obesity are known to profoundly affect HDL metabolism, composition, and subclass distribution [46], which may explain why the impact of sex becomes more pronounced in older obese populations. This pattern is consistent with prior epidemiological studies showing that menopause, BMI, and age jointly shape lipid profiles, with menopause-associated lipid changes being more evident in lean women, but less so in obese women. After identifying each subgroup, we estimated the mediation effect within each subgroup using the 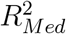 and SOS measures based on Assumption (1); the results in Figure 9. Moreover, to illustrate the relationships among the exposure (Sex), the mediator, and the outcome (HDL) within each subgroup, we used the product measure to present the coefficients derived under Assumption (1). The results are shown in the Sankey plots (Figure 10) and in Appendix Table 1-5. At the gene level, MMP8 and SQLE exhibited particularly interpretable subgroup-specific mediation patterns. For MMP8, the product-based indirect effect was most pronounced among individuals with BMI (≤ 25) and age (*>* 50) (product (= 0.0157); mediation proportion (= 4.33%)) and those with BMI (*>* 25) and age (≤ 62) (product (= 0.0129); mediation proportion (= 3.24%)), whereas the corresponding estimates were close to zero in the younger, lower-BMI subgroup and substantially attenuated in the older, higher-BMI subgroup. In subgroup 3, both the sex–MMP8 and MMP8–HDL associations were supported ((*a* = −0.1514), (*p <* 0.001); (*b* = −0.0853), (*p* = 0.0179)). This finding is biologically plausible because MMP8 can proteolytically modify apolipoprotein A-I, the major protein component of HDL, and impair its cholesterol-efflux capacity [47]. SQLE showed a complementary pattern, with its largest product estimate observed in subgroup 3 (product (= 0.0087); mediation proportion (= 2.18%)), where both component associations were evident ((*a* = −0.0626), (*p* = 0.0034); (*b* = −0.1392), (*p <* 0.001)). SQLE encodes squalene epoxidase, a key enzyme in cholesterol biosynthesis, and was included in sterol transport and metabolism pathways that were collectively associated with HDL levels in a previous pathway-wide association study [48]. These findings suggest that the molecular pathways mediating sex-related variation in HDL differ across age and adiposity profiles.

**Figure 9.**
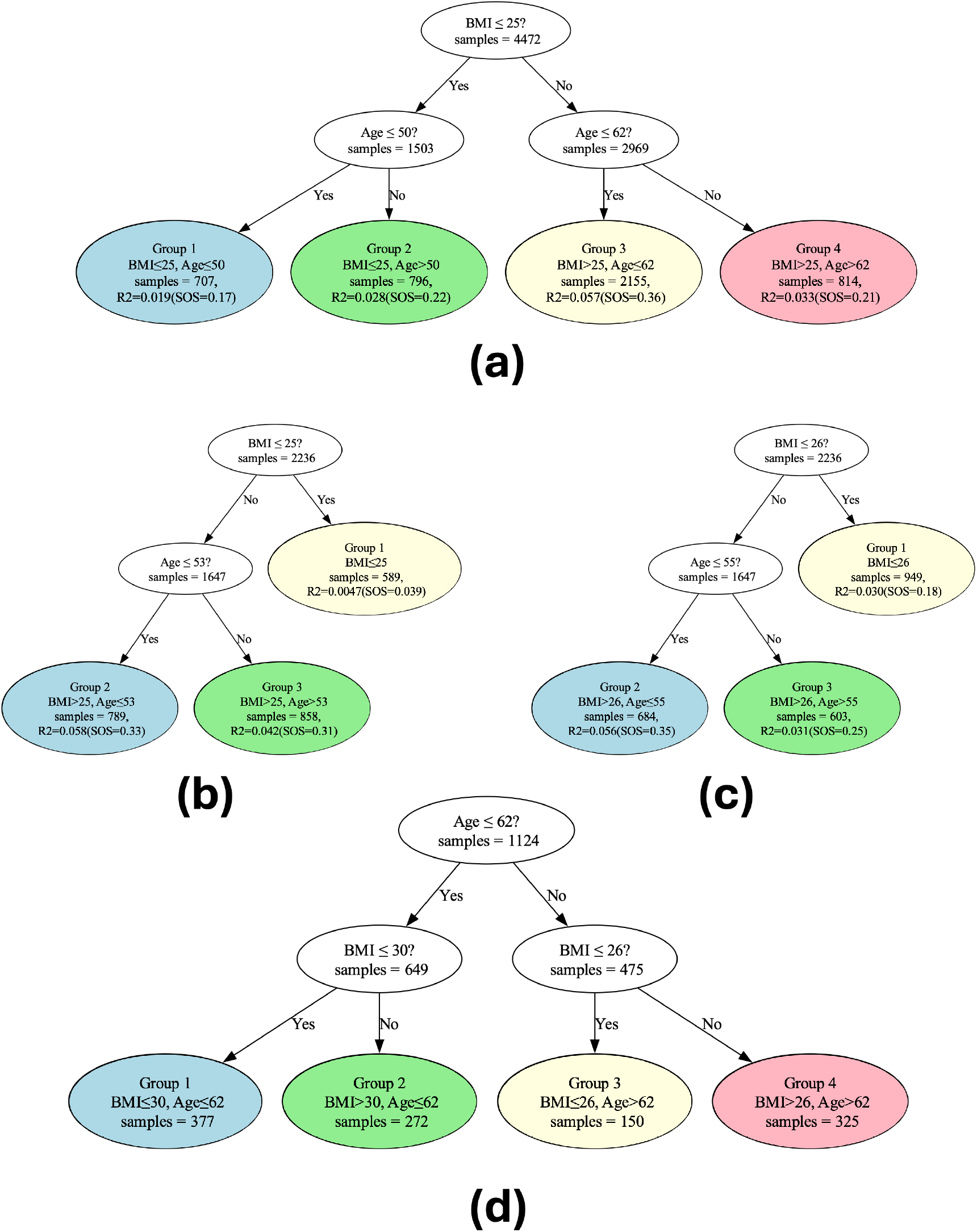
Profile for the FHS dataset and MESA dataset. (a) profile of the whole FHS dataset, (b) profile of the FHS dataset using half of the samples from random split 1, (c) profile of the FHS dataset using half of the samples from random split 2, (d) profile for the MESA dataset.

**Figure 10.**
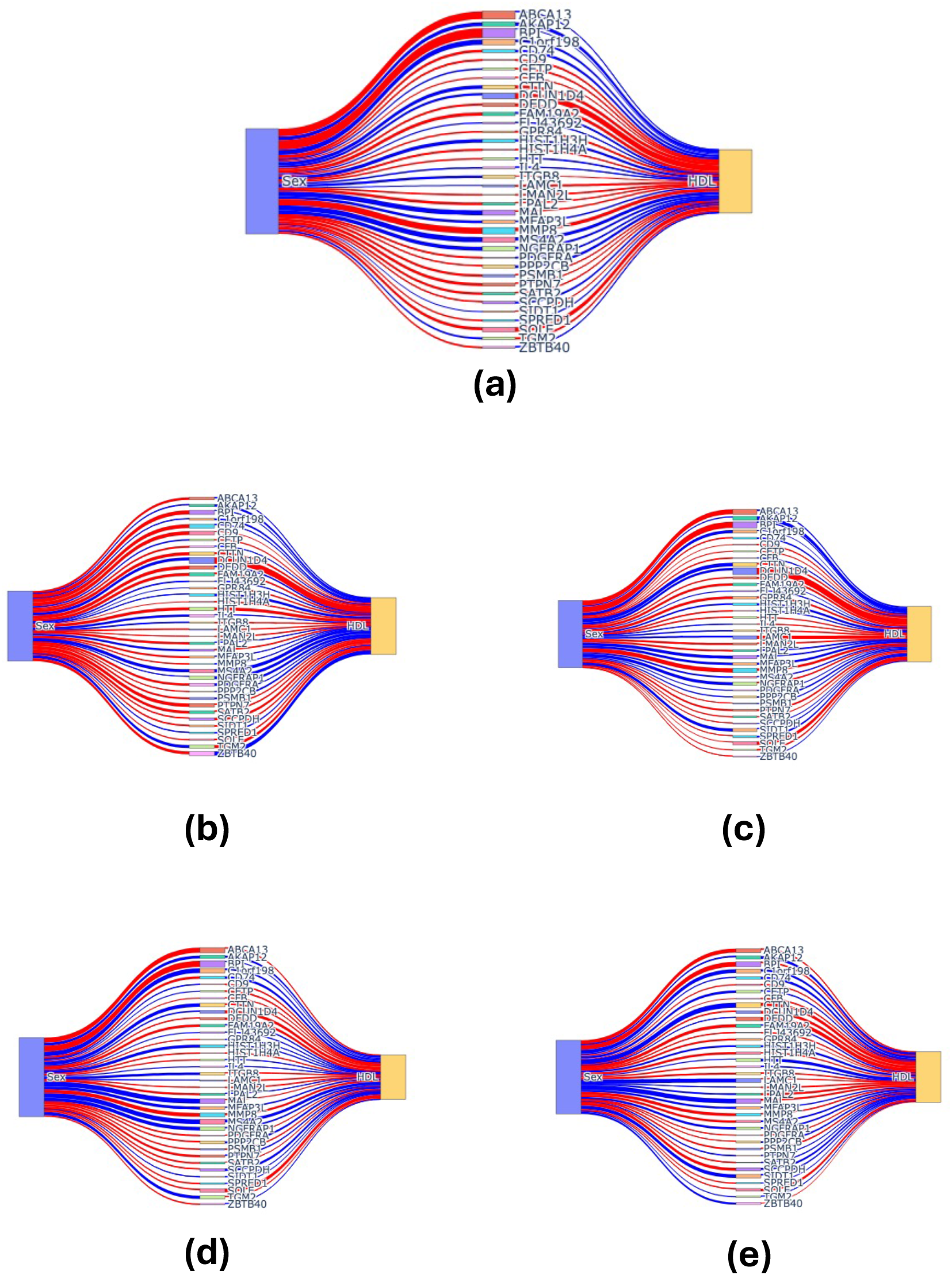
Sankey plots for the whole FHS dataset and four subgroups. Blue represents positive value, red represents negative values, width represents the line width represents the relative magnitude of the values. (a) Whole FHS dataset, (b) *BMI* ≤ 25, *Age* ≤ 50, (c)*BMI* ≤ 25, *Age >* 50, (d)*BMI >* 25, *Age* ≤ 62, (e)*BMI >* 25, *Age >* 62.

To assess robustness, we randomly split the FHS dataset into two subsets and obtained consistent results, further supporting the stability of our method (see Figure 9 (b) and (c)). Although each group had a relatively small sample size due to the partitioning of the data, the results remained consistent with those obtained from the full FHS dataset, demonstrating both the robustness of our method and the reliability of the findings.

To validate the generalizability of our findings, we also applied our method to the MESA study. The results were consistent with those from the FHS, as shown in Figure 9 (d), including selecting both BMI and age as the covariates and similar thresholds (26 and 30 for BMI corresponding to overweight and obesity and 62 years old for age). Given the smaller sample size of the MESA dataset, estimates within each subgroup were highly unstable. Accordingly, only the subgroup results are presented, without reporting specific numerical metrics.

Together, these findings demonstrate that the effect of sex on HDL through gene expression varies across subgroups defined by age and BMI. Importantly, consistent conclusions were obtained across datasets and random data partitions, further confirming the reliability of our approach and conclusions.

## Discussion

This work makes several methodological contributions to the evaluation of mediators for high dimensional settings. First, we extend the M-learner framework to accommodate high-dimensional mediators and covariates, substantially broadening its applicability in genomic and epidemiologic studies. An important strength of the proposed framework is its ability to accommodate multiple, potentially correlated mediators, a common feature of genetic and genomic studies in which molecular traits may be co-regulated or involved in shared biological pathways. In the formulation of the CAIE (5), the mediator **M** is defined as a vector rather than a single variable, allowing the joint indirect effect of correlated mediators to be characterized without requiring them to be considered in isolation. During estimation, random forests are used to predict the mediator vector, providing a flexible, data-adaptive approach that can accommodate correlations among mediators as well as nonlinear relationships and complex interactions involving the exposure and baseline covariates. The combination of the vector-valued CAIE formulation and random-forest-based estimation therefore makes the proposed method particularly well suited to mediation analyses involving multiple correlated molecular features. Second, compared with low-dimensional settings, the definition of the *p*_leaf_ metric differs from that in [11].

In high-dimensional contexts, the original metric proposed in [11] may become unstable or fail to adequately capture subgroup structure; accordingly, we introduce a modified formulation tailored to high-dimensional settings.

Taken together, these innovations provide a flexible, data-adaptive framework for investigating heterogeneity in mediators and for understanding when specific molecular mediators are more or less informative about exposure effects in high-dimensional settings, particularly in omics studies. By enabling the identification of context-dependent pathways and subgroup-specific mechanisms, this framework offers a principled approach for improving biological interpretability and advancing the study of complex molecular systems.

In high-dimensional genomic settings, the effects of molecular mediators are unlikely to be uniform across the population, as regulatory pathways may contribute differently to disease processes under varying genetic and environmental conditions. Accounting for heterogeneity in mediation effects is therefore critical for accurately characterizing the functional roles of candidate mediators and for identifying subpopulations in which specific molecular mechanisms drive phenotypic variation. Ignoring such heterogeneity may lead to incomplete or misleading interpretations of genomic data, particularly when signals are present only in a subset of individuals. Methodological frameworks that explicitly model heterogeneous mediation effects can thus improve biological interpretability and enhance the utility of genomic analyses for understanding complex disease biology.

The central idea of our approach is to construct similarity based on CAIE. Specifically, we begin with a exposure distance matrix (i.e., a dissimilarity matrix) derived from estimated CAIE. A modified t-SNE procedure is then used to transform dissimilarities into similarities and embed observations into a low-dimensional Euclidean space. This representation facilitates the application of the K-means algorithm to identify subgroups with homogeneous indirect effects.

Consistent with prior work [11], t-SNE demonstrates superior performance relative to UMAP across diverse scenarios, whereas UMAP retains advantages in computational efficiency and scalability to very high-dimensional settings [49]. Simulation studies indicate that the proposed approach remains robust under high-noise conditions, effectively controls type I error, and reliably detects heterogeneity, underscoring its potential utility for elucidating complex biological mechanisms and informing precision intervention strategies [11].

Our framework is also robust to the specification of the maximum number of clusters (*K*), with results remaining stable across a reasonable range of candidate values. To enhance interpretability, we summarize clustering results using decision trees defined by terminal nodes rather than relying directly on raw K-means assignments. This strategy ensures that each subgroup is characterized by distinct covariate profiles while reducing the risk of over-fitting. Additional structural constraints, such as limiting each variable to a single split per branch and restricting minimum leaf size and maximum tree depth, further promote interpretability without materially compromising model fidelity [11].

The application of the proposed method to the FHS dataset demonstrates its practical utility in uncovering biologically meaningful heterogeneity in mediation effects within complex population data. The identified subgroups align with well-established clinical and physiological patterns linking age, BMI, and sex-specific hormonal changes to HDL metabolism, providing external support for the validity of the detected subgroup structure. In particular, the method was able to distinguish context-dependent mechanisms that are consistent with known differences in lipid regulation across menopausal status and obesity levels.

These findings suggest that the proposed framework is capable of capturing subtle yet clinically relevant variation in molecular or physiological pathways that may be overlooked by conventional approaches assuming homogeneous mediation effects. More broadly, the results illustrate the potential of the proposed method to generate interpretable subgtype-specific insights in real-world genomic and epidemiologic studies.

Several limitations warrant consideration. First, the current framework is limited to binary exposures; extensions to continuous exposures and survival outcomes would enable more nuanced characterization of exposure–outcome relationships and broaden its applicability. Second, mediators are initially screened using the full sample based on 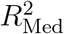 –based procedure, after which heterogeneity is assessed separately for each selected mediator and mediators showing evidence of heterogeneity are jointly analyzed. Although this two-stage strategy may miss some heterogeneous mediators, jointly evaluating all candidates is often infeasible in studies with limited sample sizes; thus, our approach deliberately prioritizes finite-sample stability and reliability over maximal sensitivity. Finally, because the method is designed primarily to identify heterogeneous mediators, homogeneous mediators may receive less attention. Developing a unified framework that simultaneously accommodates homogeneous and heterogeneous mediators remains an important direction for future research.

## Supporting information

Main manuscript

## Data Availability and Implementation

The proposed method is available at https://github.com/1996lixingyu1996/M-high-lear The data sets used for the analyses described in this manuscript were obtained from NIH/dbGaP at https://www.ncbi.nlm.nih.gov/gap/ through accession numbers phs000007 phs000974, phs000209, and phs001416.

## Funding

This research was supported by National Institutes of Health (NIH) grants R21HL170213 and R01HL184065. The content is solely the responsibility of the authors and does not necessarily represent the official views of the NIH. The Framinham Heart Study (FHS) is conducted and supported by the National Heart, Lung, and Blood Institute (NHLBI) in collaboration with Boston University. The Multi-Ethnic Study of Atherosclerosis (MESA) is conducted and supported by the NHLBI in collaboration with MESA investigators. This manuscript was not prepared in collaboration with investigators in the FHS or MESA and does not necessarily reflect the opinions or views of the FHS, Boston University, the MESA, or the NHLBI.

## Competing interests

The authors have declared that no competing interests exist.

## Supporting information

The Supporting Information includes the following sections:

**Appendix A**: Details on the t-SNE

**Appendix B**: Pipeline of the M-learner

**Appendix C**: Additional simulations and results

**Appendix D**: Supplemental information of the real data applications

## Notes

https://github.com/1996lixingyu1996/M-high-learner

### Competing Interest Statement

The authors have declared no competing interest.

