## Supplementary material for "An M-learner approach for heterogeneous mediation analysis with high-dimensional omics mediators": Main manuscript

**Appendix A**: Details on the t-SNE

**Appendix B**: Pipeline of the M-learner

**Appendix C**: Additional simulations and results

**Appendix D**: Supplemental information of the real data applications

### Appendix A: Details on the t-SNE

Here, we introduce the process of modified t-SNE projection in our method. To simplify the notation, we denote $dis(i,j)$ by $d_{ij}$.
In modofoed t-SNE projection process, the first step is to calculate the similarity probability:

$$p_{j|i} = \frac{\exp\left(-d_{ij}^2/2\sigma_i^2\right)}{\sum_{k\neq i}\exp(-d_{ik}^2/2\sigma_i^2)} \tag{1}$$

where $p_{j|i}$ is the similarity probability, $\sigma_i$ is determined by preplexity, which is chosen by user, the detailed introduction can be found in [Maaten and Hinton, 2008, Cai and Ma, 2022]. In original t-SNE method, the distance is obtained by calculated the distance of $i$ and $j$. In our method, we use indirect effect distance to replace the original distance.
Then symmetric similarity probability

$$p_{ij} = \frac{p_{j|i} + p_{i|j}}{2n},$$

Then, defined the similarity in low dimension (Euclidean space),

$$q_{ij} = \frac{(1 + \|o_i - o_j\|^2)^{-1}}{\sum_{k\neq i}\exp(1 + \|o_k - o_l\|^2)^{-1}},$$

where $o_i$ represents the coordinates of the $i$-th individual in the Euclidean embedding.
Finally, t-SNE optimizes the Kullback–Leibler divergence

$$\min \sum_{i\neq j} p_{ij} \log \frac{p_{ij}}{q_{ij}}$$

to determine the coordinates of different units in the Euclidean space.
The theoretical analysis is provided in Theorems S2 and S3 of [Li et al., 2026].

### Appendix B: Pipeline of the M-learner

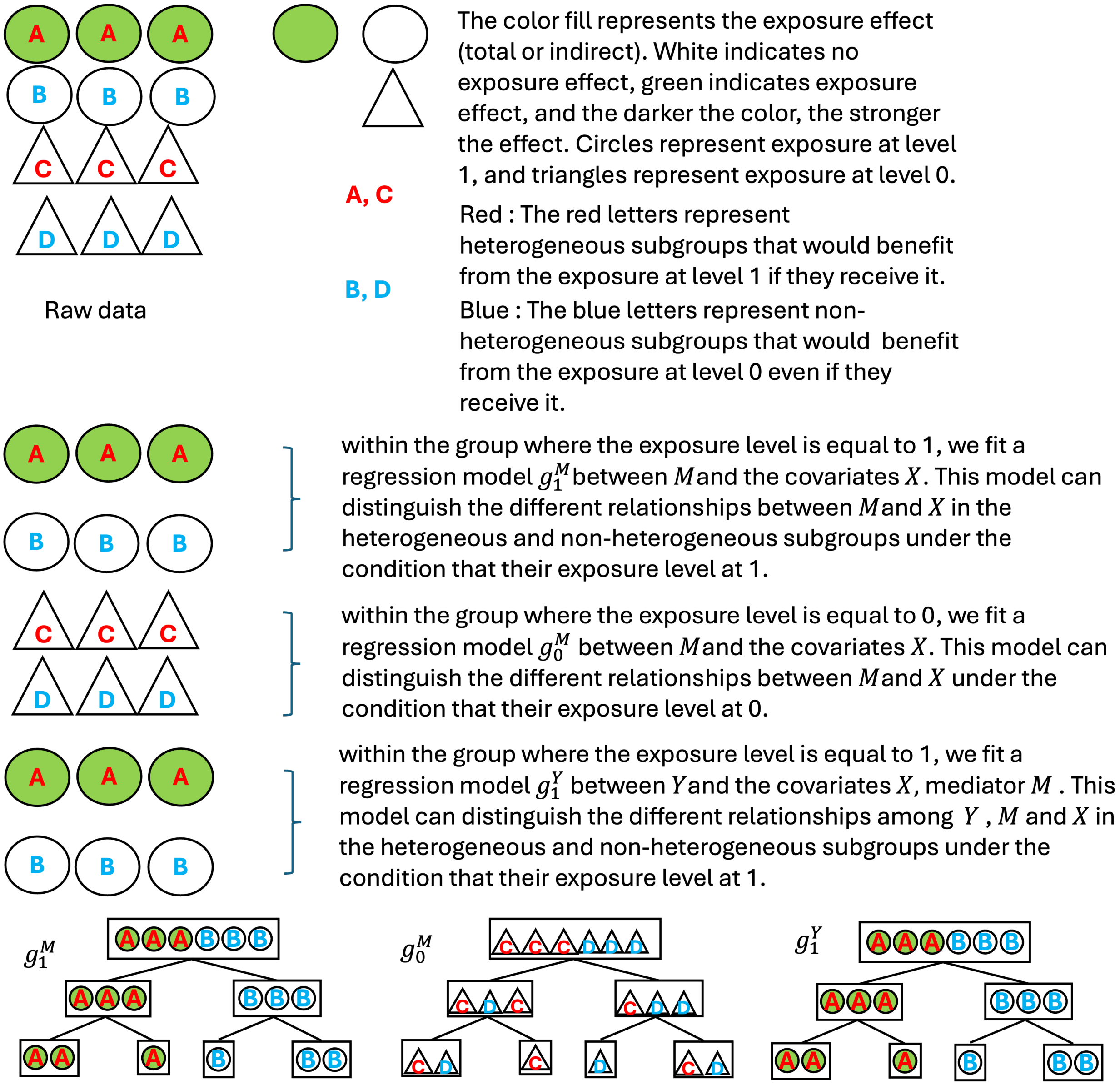


**Figure 1.** The pipeline of the M-high-learner method (A). This flowchart illustrates the data and the three models.

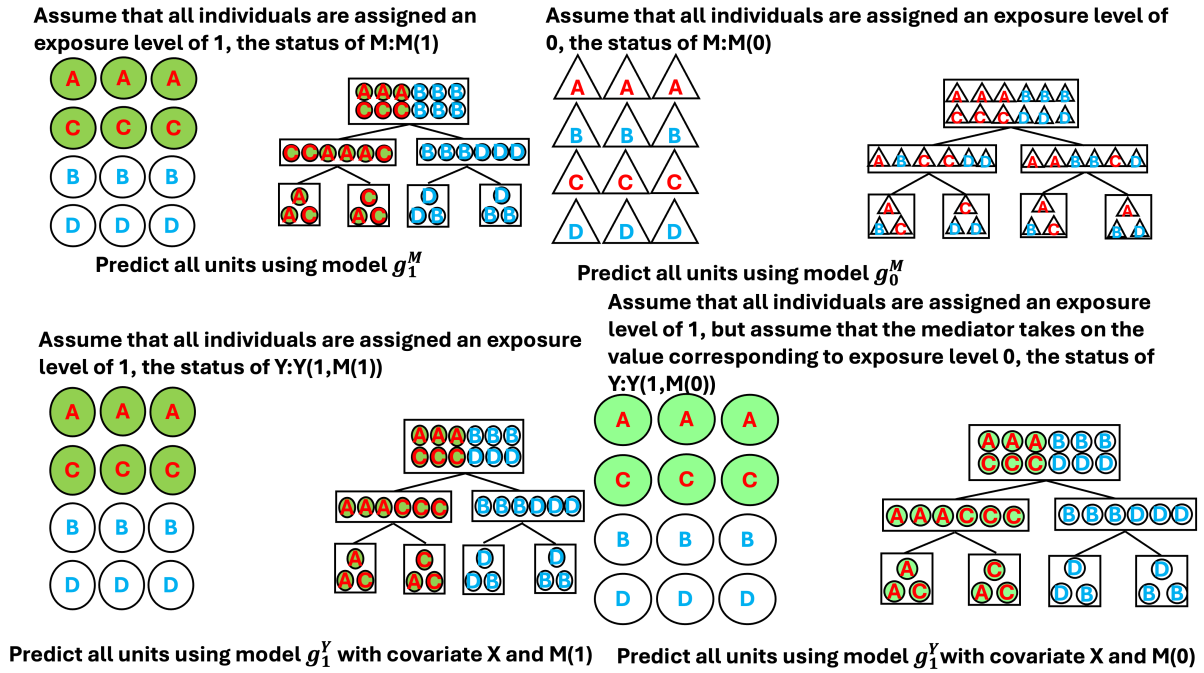


**Figure 2.** The pipeline of the M-high-learner method (B). In this part of the flowchart, we demonstrate how to compute the CAIE.

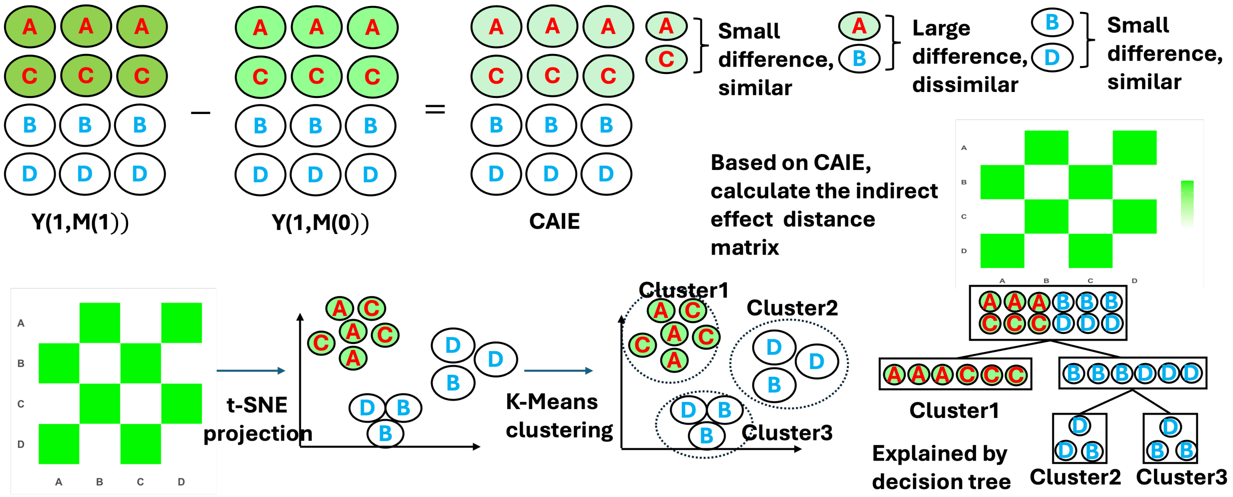


**Figure 3.** The pipeline of the M-high-learner method (C). This flowchart illustrates how to compute the treatment distance matrix, perform t-SNE projection and clustering, and use a decision tree for interpretation.

#### Decision-tree constraints

To improve subgroup interpretability and reduce the risk of overfitting, three constraints may be imposed during decision-tree construction. First, the minimum leaf-size constraint requires each terminal node, and hence each identified subgroup, to contain at least a prespecified number of individuals. This prevents the identification of very small, data-driven subgroups for which effect estimates may be unstable. Second, the maximum-depth constraint limits the number of successive splits from the root node to any terminal node, thereby controlling tree complexity. Third, the non-repeated splitting-variable constraint allows each covariate to be used at most once within a tree. This restriction avoids redundant splits based on multiple cutoffs of the same variable and produces simpler, more interpretable subgroup definitions.
In the present study, we applied only the minimum leaf-size and non-repeated splitting-variable constraints. We did not impose a prespecified maximum tree depth; instead, tree complexity was controlled indirectly through the other two restrictions. The maximum depth of 2 shown in Web Figure 4(a) is therefore provided solely to illustrate this optional constraint.

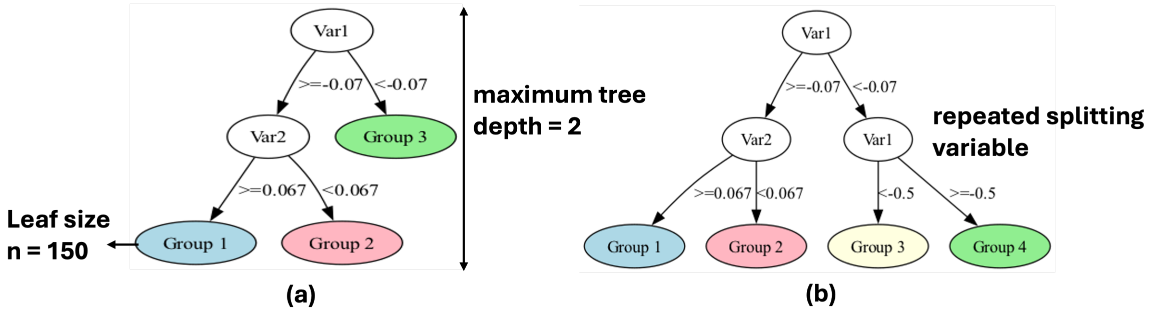


**Figure 4.** Illustration of potential constraints on the decision tree used to explain the identified clusters. (a) A decision tree illustrating three possible constraints: a minimum leaf size of 150, a maximum depth of 2, and no repeated splitting variables. (b) A tree that violates the non-repeated splitting-variable constraint because Var1 is used at both the root node and a subsequent internal node. In the present study, only the minimum leaf-size and non-repeated splitting-variable constraints were imposed; the maximum-depth constraint is shown for illustrative purposes only.

### Appendix C: Additional simulations and results

#### Single-Mediator

All functions were defined for units under exposure level at ($w = 1$) and level at ($w = 0$), and in all scenarios the treatment probability is assigned with equation (2),

$$W \mid X = \mathbf{x} \sim \text{Bernoulli}\{0.5)\}, \tag{2}$$

$$
\begin{aligned}
M(w) &= \eta^{(1)}(\mathbf{x}) + \frac{1}{2}(2w-1)\cdot\kappa^{(1)}(\mathbf{x}) + b^{(1)} + \epsilon^{(1)}, \\
Y(w) &= \eta^{(2)}(\mathbf{x}) + \frac{1}{2}(2w-1)\cdot\kappa^{(2)}(\mathbf{x}) + b^{(2)} + c\cdot M(w) + \epsilon^{(2)}, \qquad (3)
\end{aligned}
$$

where $\epsilon^{(1)} \sim \mathcal{N}(0, 0.01)$, $\epsilon^{(2)} \sim \mathcal{N}(0, 0.01)$, and the $\mathbf{x}$ are independent of $\epsilon^{(1)}$, $\epsilon^{(2)}$ and one another, and $x^{(j)} \sim \mathcal{N}(0,1)$,for $j = 1, 2, \cdots, 50$, $b^{(1)}$ and $b^{(2)}$ are the intercept terms, $c$ is the coefficient of mediator.

1. In (3), existing heterogeneity, all exposure effects via mediator (All):$\eta^{(1)}(\mathbf{x}) = \frac{1}{2}(x^{(1)} + x^{(2)}) + x^{(3)} + x^{(4)}$,$\kappa_1(\mathbf{x}) = \sum_{i=1}^{2}\mathbb{I}(x^{(i)} > 0)\cdot x^{(i)}$, $b^{(1)} = 0$, $\eta^{(2)}(\mathbf{x}) = \frac{1}{2}(x^{(1)} + x^{(2)}) + x^{(3)} + x^{(4)})$,$\kappa^{(2)}(x) = 0$, $b^{(2)} = 1, c = 1$.

2. In (3), existing heterogeneity, part of the exposure effect via mediator (Part): $\eta^{(1)}(\mathbf{x}) = \frac{1}{2}(x^{(1)} + x^{(2)}) + x^{(3)} + x^{(4)})$,$\kappa^{(1)}(\mathbf{x}) = \sum_{i=1}^{2}\mathbb{I}(x^{(i)} > 0)\cdot x^{(i)}$, $b^{(1)} = 0, c = 1$, $\eta^{(2)}(\mathbf{x}) = \frac{1}{2}(x^{(1)} + x^{(2)}) + x^{(3)} + x^{(4)})$,$\kappa^{(2)}(\mathbf{x}) = \sum_{i=1}^{2}\mathbb{I}(x^{(i)} > 0)\cdot x^{(i)}$, $b^{(2)} = 1, c = 1$.

3. In (3), no heterogeneity, all units benefit from the exposure (Global):$\eta^{(1)}(\mathbf{x}) = \frac{1}{2}(x^{(1)} + x^{(2)}) + x^{(3)} + x^{(4)})$,$\kappa^{(1)}(\mathbf{x}) = 1$, $b^{(1)} = 0$, $\eta^{(2)}(\mathbf{x}) = \frac{1}{2}(x^{(3)} + x^{(4)})$,$\kappa^{(2)}(x) = 0$, $b^{(2)} = 1, c = 1$.

4. In (3), no heterogeneity, 0% exposure effects via mediator(Null 1):$\eta^{(1)}(\mathbf{x}) = \frac{1}{2}(x^{(1)}+x^{(2)}) + x^{(3)} + x^{(4)})$,$\kappa^{(1)}(x) = 0$, $b^{(1)} = 0$, $\eta^{(2)}(\mathbf{x}) = \frac{1}{2}(x^{(3)} + x^{(4)})$,$\kappa_2(\mathbf{x}) = \sum_{i=1}^{2}\mathbb{I}(x^{(i)} > 0)\cdot x^{(i)}$, $b^{(2)} = 1, c = 1$.

5. In (3), no heterogeneity, 0% exposure effects via mediator(Null 2):$\eta^{(1)}(\mathbf{x}) = \frac{1}{2}(x^{(1)} + x^{(2)}) + x^{(3)} + x^{(4)})$,$\kappa^{(1)}(\mathbf{x}) = \sum_{i=1}^{2}\mathbb{I}(x^{(i)} > 0)\cdot x^{(i)}$, $b^{(1)} = 0$, $\eta^{(2)}(\mathbf{x}) = \frac{1}{2}(x^{(3)} + x^{(4)})$,$\kappa^{(2)}(\mathbf{x}) = \sum_{i=1}^{2}\mathbb{I}(x^{(i)} > 0)\cdot x^{(i)}$, $b^{(2)} = 1, c = 0$.

6-10. Compared to Setting 1, the only change is the sample size, which is increased to 2,000. All other experimental conditions are held constant.

Across all simulation experiments, the number of trees in the random forest was fixed at 2000, while all other parameters were kept at their default settings. All analyses were implemented in R (version 4.3.3) using the `randomForestSRC` package (version 3.2.3).

**Multiple-Mediators**

All functions were defined for units under exposure at level ($w = 1$) and level at ($w = 0$), and in all scenarios the exposure probability is assigned with equation,
All functions were defined for units under exposure at level ($w = 1$) and level ($w = 0$), and

in all scenarios the exposure probability is assigned with equation (4),

$$
\begin{aligned}
W \mid X = \mathbf{x} &\sim \text{Bernoulli}\{0.5\}, \\
M^{(j)}(w) &= \eta^{(1j)}(\mathbf{x}) + \frac{1}{2}(2w-1)\,\kappa^{(1)}(\mathbf{x}) + b^{(1)} + \epsilon^{(1j)}, \quad j = 1, \ldots, 10, \\
Y(w) &= \eta^{(2)}(\mathbf{x}) + \frac{1}{2}(2w-1)\,\kappa^{(2)}(\mathbf{x}) + b^{(2)} + c\sum_{j=1}^{10} M^{(j)}(w) + \epsilon^{(2)}.
\end{aligned} \tag{4}
$$

where $\epsilon^{(1j)} \sim \mathcal{N}(0, 0.01)$, $\epsilon^{(2)} \sim \mathcal{N}(0, 0.01)$, and the $\mathbf{x}$ are independent of $\epsilon^{(1j)}$, $\epsilon^{(2)}$ and one another, and $x^{(i)} \sim \mathcal{N}(0,1)$, $i = 1, \cdots, 50$, $b^{(1)}$ and $b^{(2)}$ are the intercept terms, $c$ is the coefficient of the mediator.

11. In (4), existing heterogeneity, part of the exposure effect via mediator (All): $\eta^{(1j)}(\mathbf{x}) = \sum_{i=1}^{4} a^{(ji)} x^{(i)}$, where

$$
\begin{aligned}
a^{(j1)} &= 0.5 + (1/30)(j-1) - (0.15), \\
a^{(j2)} &= 0.5 - (1/30)(j-1) + (0.15), \\
a^{(j3)} &= 1 + (1/30)(j-1) - (0.15), \\
a^{(j4)} &= 1 - (1/30)(j-1) + (0.15),
\end{aligned}
$$

$\kappa^{(1)}(\mathbf{x}) = \sum_{i=1}^{2} \mathbb{I}(x^{(i)} > 0) \cdot x_i$, $b_1 = 0$, $\eta^{(2)}(\mathbf{x}) = \frac{1}{2}(x^{(1)} + x^{(2)}) + x^{(3)} + x^{(4)}$, $\kappa^{(2)}(\mathbf{x}) = 0$, $b^{(2)} = 1, c = 1$, $\kappa^{(2)}(\mathbf{x}) = \sum_{i=1}^{2} \mathbb{I}(x^{(i)} > 0) \cdot x^{(i)}$, $b^{(2)} = 1, c = 0.1$.

12. In (4), all setting is the same with 11, except $\kappa^{(2)}(x) = \sum_{i=1}^{2} \mathbb{I}(x^{(i)} > 0) \cdot x^{(i)}$.

13. In (4), no heterogeneity, all units benefit from the exposure at level 1 (Global): all setting is the same with 11, except $\kappa^{(2)}(\mathbf{x}) = 1$.

14. In (4), no heterogeneity, 0% treatment effects via mediator (Null 1): $\eta^{(1)}(\mathbf{x}) = \frac{1}{2}(x^{(1)} + x^{(2)} + x^{(3)} + x^{(4)}$,$\kappa_1(\mathbf{x}) = 0$, $b_1 = 0$, $\eta_2(\mathbf{x}) = \frac{1}{2}(x^{(3)} + x^{(4)})$,

    $\kappa_2(\mathbf{x}) = \sum_{i=1}^{2} \mathbb{I}(x^{(i)} > 0) \cdot x^{(i)}$, $b^{(2)} = 1, c = 0.1$.

15. In (4), no heterogeneity, 0% exposure effects via mediator (Null 2):all setting is the same with 11, except $c = 0$

16-20. Compared to Setting 11-15, the only change is the sample size, which is increased to $2,000$. All other experimental conditions are held constant.

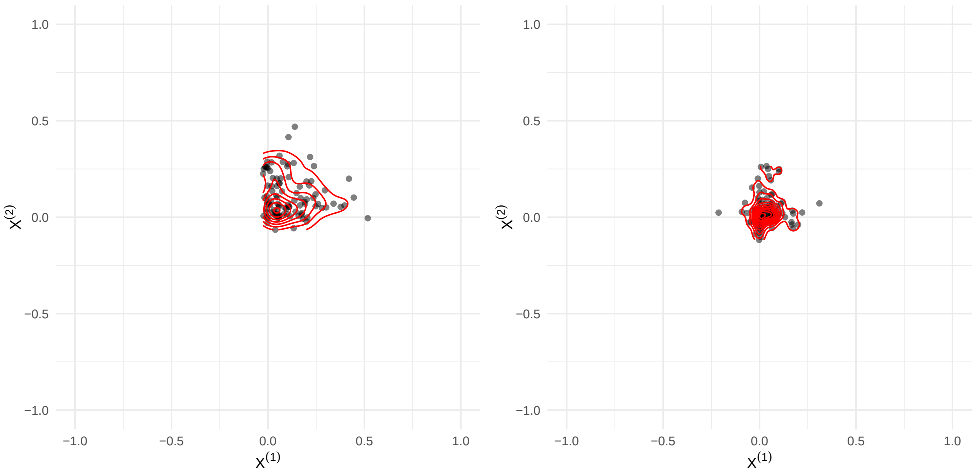


**Figure 5.** Threshold distributions for single mediator when sample size is 2000. Solid dots represent the estimated thresholds of heterogeneous subgroups obtained in each experiment, with values greater than 1 or undefined thresholds set to 1. The left panel is All scenario, the right panel is Part scenario.

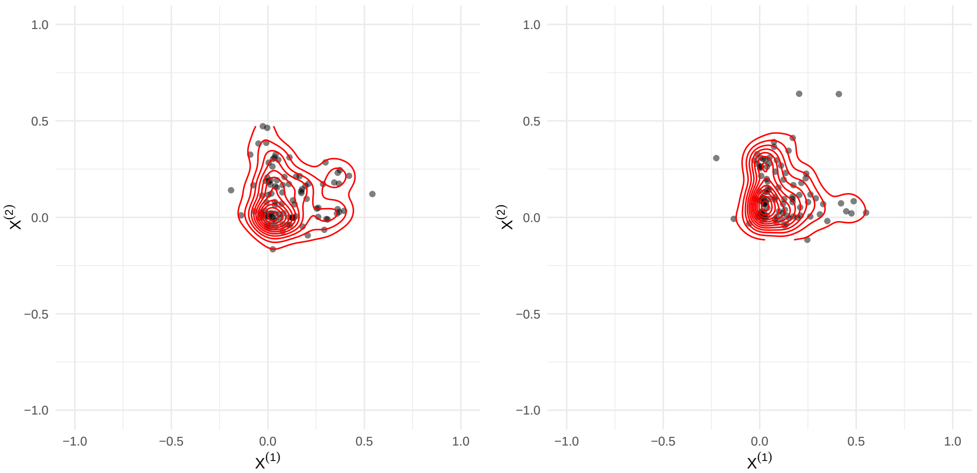


**Figure 6.** Threshold distributions for multiple mediators when sample size is 1000. Solid dots represent the estimated thresholds of heterogeneous subgroups obtained in each experiment, with values greater than 1 or undefined thresholds set to 1. The left panel is All scenario, the right panel is Part scenario.

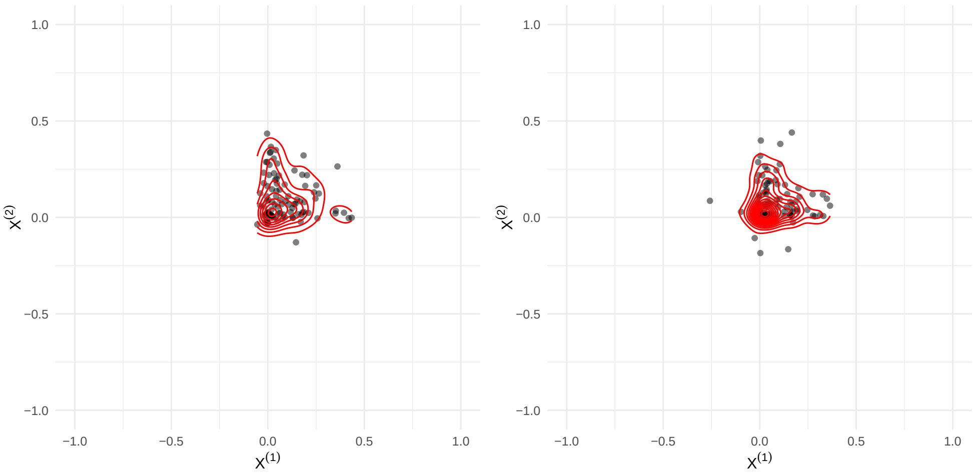


**Figure 7.** Threshold distributions for multiple mediators when sample size is 2000. Solid dots represent the estimated thresholds of heterogeneous subgroups obtained in each experiment, with values greater than 1 or undefined thresholds set to 1. The left panel is All scenario, the right panel is Part scenario.

### Appendix D: Supplemental information of the real data applications

| Mediator | a | p.value(a) | b | p.value(b) | Product | Med Prop |
|---|---|---|---|---|---|---|
| ZBTB40 | -0.0534 | 0.0003 | 0.0140 | 0.5876 | -0.0007 | -0.0018 |
| LAMC1 | 0.0321 | 0.0303 | -0.0592 | 0.0001 | -0.0019 | -0.0045 |
| DEDD | -0.0626 | 0.0000 | -0.0901 | 0.0000 | 0.0056 | 0.0132 |
| PTPN7 | -0.0839 | 0.0000 | -0.0020 | 0.9193 | 0.0002 | 0.0004 |
| MAL | 0.1362 | 0.0000 | -0.0431 | 0.0081 | -0.0059 | -0.0138 |
| SATB2 | -0.0660 | 0.0000 | -0.0141 | 0.3164 | 0.0009 | 0.0022 |
| SIDT1 | 0.0342 | 0.0210 | -0.0053 | 0.7837 | -0.0002 | -0.0004 |
| HTT | -0.0515 | 0.0005 | 0.0164 | 0.5551 | -0.0008 | -0.0020 |
| IL4 | 0.0479 | 0.0012 | 0.0064 | 0.6446 | 0.0003 | 0.0007 |
| CD74 | -0.0755 | 0.0000 | 0.0811 | 0.0001 | -0.0061 | -0.0144 |
| HIST1H3H | 0.0965 | 0.0000 | 0.0723 | 0.0000 | 0.0070 | 0.0164 |
| CFB | -0.0426 | 0.0040 | -0.0027 | 0.8604 | 0.0001 | 0.0003 |
| AKAP12 | 0.1089 | 0.0000 | 0.1029 | 0.0000 | 0.0112 | 0.0263 |
| LPAL2 | -0.0734 | 0.0000 | -0.0104 | 0.4349 | 0.0008 | 0.0018 |
| PSMB1 | -0.0648 | 0.0000 | -0.0109 | 0.6322 | 0.0007 | 0.0017 |
| ITGB8 | 0.0737 | 0.0000 | -0.0350 | 0.0311 | -0.0026 | -0.0060 |
| MS4A2 | 0.1376 | 0.0000 | 0.0933 | 0.0000 | 0.0128 | 0.0301 |
| SCCPDH | -0.0150 | 0.3122 | 0.0671 | 0.0018 | -0.0010 | -0.0024 |
| C1orf198 | 0.1462 | 0.0000 | 0.0691 | 0.0002 | 0.0101 | 0.0237 |
| LMAN2L | -0.0166 | 0.2635 | -0.0180 | 0.1974 | 0.0003 | 0.0007 |
| DCUN1D4 | 0.0619 | 0.0000 | -0.1637 | 0.0000 | -0.0101 | -0.0238 |
| PDGFRA | -0.0241 | 0.1041 | 0.0231 | 0.1804 | -0.0006 | -0.0013 |
| MFAP3L | 0.0781 | 0.0000 | 0.0460 | 0.0162 | 0.0036 | 0.0084 |
| HIST1H4A | -0.0296 | 0.0458 | -0.0105 | 0.4222 | 0.0003 | 0.0007 |
| ABCA13 | -0.2173 | 0.0000 | 0.0062 | 0.7995 | -0.0013 | -0.0031 |
| FLJ43692 | 0.0382 | 0.0100 | 0.0074 | 0.5876 | 0.0003 | 0.0007 |
| SQLE | -0.0799 | 0.0000 | -0.1241 | 0.0000 | 0.0099 | 0.0232 |
| PPP2CB | -0.0142 | 0.3382 | 0.0733 | 0.0001 | -0.0010 | -0.0024 |
| CTTN | 0.0996 | 0.0000 | -0.0897 | 0.0000 | -0.0089 | -0.0209 |
| MMP8 | -0.1806 | 0.0000 | -0.1015 | 0.0000 | 0.0183 | 0.0430 |
| CD9 | -0.0289 | 0.0511 | -0.0271 | 0.0866 | 0.0008 | 0.0018 |
| GPR84 | -0.0458 | 0.0020 | -0.0288 | 0.0425 | 0.0013 | 0.0031 |
| FAM19A2 | -0.0909 | 0.0000 | 0.0288 | 0.0383 | -0.0026 | -0.0061 |
| SPRED1 | -0.0494 | 0.0009 | 0.0431 | 0.0031 | -0.0021 | -0.0050 |
| CETP | 0.0450 | 0.0024 | -0.0244 | 0.0999 | -0.0011 | -0.0026 |
| BPI | -0.2606 | 0.0000 | 0.0307 | 0.1680 | -0.0080 | -0.0187 |
| TGM2 | 0.0543 | 0.0002 | -0.0600 | 0.0002 | -0.0033 | -0.0076 |
| NGFRAP1 | 0.1135 | 0.0000 | 0.0639 | 0.0010 | 0.0073 | 0.0170 |
| Direct Effect | - | - | - | - | 0.3932 | 0.0000 |

**Table 1.** Product measure estimates in FHS dataset. For Direct effect, the product is the estimation of indirect effect. Med Prop: Mediation proportion.

| Mediator | a | p.value(a) | b | p.value(b) | product | med prop |
|---|---|---|---|---|---|---|
| ZBTB40 | -0.1108 | 0.0029 | 0.1619 | 0.0201 | -0.0179 | -0.0533 |
| LAMC1 | -0.0260 | 0.4857 | -0.0523 | 0.2560 | 0.0014 | 0.0040 |
| DEDD | -0.1193 | 0.0013 | 0.0287 | 0.5337 | -0.0034 | -0.0102 |
| PTPN7 | -0.1263 | 0.0007 | -0.0535 | 0.3085 | 0.0068 | 0.0201 |
| MAL | 0.0775 | 0.0373 | -0.0259 | 0.5560 | -0.0020 | -0.0060 |
| SATB2 | -0.1007 | 0.0068 | -0.0290 | 0.4766 | 0.0029 | 0.0087 |
| SIDT1 | -0.0288 | 0.4399 | -0.0635 | 0.2250 | 0.0018 | 0.0054 |
| HTT | -0.1117 | 0.0027 | -0.1140 | 0.1628 | 0.0127 | 0.0379 |
| IL4 | 0.0878 | 0.0182 | 0.0081 | 0.8322 | 0.0007 | 0.0021 |
| CD74 | -0.1382 | 0.0002 | 0.0093 | 0.8746 | -0.0013 | -0.0038 |
| HIST1H3H | 0.0106 | 0.7760 | 0.0940 | 0.0399 | 0.0010 | 0.0030 |
| CFB | -0.0648 | 0.0816 | 0.0059 | 0.8877 | -0.0004 | -0.0011 |
| AKAP12 | 0.0222 | 0.5517 | 0.0845 | 0.0852 | 0.0019 | 0.0056 |
| LPAL2 | -0.0875 | 0.0187 | 0.0086 | 0.8172 | -0.0008 | -0.0022 |
| PSMB1 | -0.0731 | 0.0497 | 0.0720 | 0.2775 | -0.0053 | -0.0156 |
| ITGB8 | 0.0266 | 0.4748 | -0.0495 | 0.2483 | -0.0013 | -0.0039 |
| MS4A2 | 0.1178 | 0.0015 | 0.1236 | 0.0121 | 0.0146 | 0.0433 |
| SCCPDH | -0.0233 | 0.5317 | -0.0957 | 0.1311 | 0.0022 | 0.0066 |
| C1orf198 | 0.0180 | 0.6296 | 0.0701 | 0.1557 | 0.0013 | 0.0037 |
| LMAN2L | -0.0031 | 0.9330 | -0.0183 | 0.6392 | 0.0001 | 0.0002 |
| DCUN1D4 | 0.1062 | 0.0043 | -0.2327 | 0.0025 | -0.0247 | -0.0735 |
| PDGFRA | -0.0900 | 0.0156 | 0.0956 | 0.0377 | -0.0086 | -0.0256 |
| MFAP3L | -0.0693 | 0.0629 | 0.0071 | 0.8844 | -0.0005 | -0.0015 |
| HIST1H4A | -0.0372 | 0.3183 | -0.0208 | 0.5701 | 0.0008 | 0.0023 |
| ABCA13 | -0.0925 | 0.0129 | 0.0172 | 0.7650 | -0.0016 | -0.0047 |
| FLJ43692 | 0.0251 | 0.5004 | 0.0147 | 0.7043 | 0.0004 | 0.0011 |
| SQLE | -0.0481 | 0.1964 | -0.0302 | 0.4623 | 0.0015 | 0.0043 |
| PPP2CB | -0.0133 | 0.7221 | 0.0088 | 0.8688 | -0.0001 | -0.0003 |
| CTTN | -0.1208 | 0.0011 | -0.0710 | 0.2134 | 0.0086 | 0.0255 |
| MMP8 | -0.0165 | 0.6582 | 0.0097 | 0.8680 | -0.0002 | -0.0005 |
| CD9 | -0.1400 | 0.0002 | -0.0337 | 0.4522 | 0.0047 | 0.0140 |
| GPR84 | 0.0373 | 0.3170 | -0.0606 | 0.1165 | -0.0023 | -0.0067 |
| FAM19A2 | -0.1096 | 0.0032 | -0.0311 | 0.4154 | 0.0034 | 0.0101 |
| SPRED1 | 0.0168 | 0.6527 | 0.0556 | 0.1664 | 0.0009 | 0.0028 |
| CETP | 0.0588 | 0.1146 | -0.0466 | 0.2614 | -0.0027 | -0.0081 |
| BPI | -0.1563 | 0.0000 | 0.0074 | 0.8870 | -0.0012 | -0.0035 |
| TGM2 | 0.1117 | 0.0027 | -0.1082 | 0.0147 | -0.0121 | -0.0359 |
| NGFRAP1 | 0.0070 | 0.8502 | 0.1122 | 0.0447 | 0.0008 | 0.0024 |
| Direct Effect | - | - | - | - | 0.3542 | 0.0000 |

**Table 2.** Product measure estimates in FHS dataset, subgroup 1 (BMI $\leq$ 25, Age $\leq$ 50). For Direct effect, the product is the estimation of indirect effect. Med Prop: Mediation proportion.

| Mediator | a | p.value(a) | b | p.value(b) | product | med prop |
|---|---|---|---|---|---|---|
| ZBTB40 | -0.0053 | 0.8809 | 0.0417 | 0.5338 | -0.0002 | -0.0006 |
| LAMC1 | 0.0728 | 0.0377 | -0.1288 | 0.0009 | -0.0094 | -0.0258 |
| DEDD | -0.0560 | 0.1101 | -0.0835 | 0.0583 | 0.0047 | 0.0129 |
| PTPN7 | -0.0710 | 0.0428 | -0.0715 | 0.1425 | 0.0051 | 0.0140 |
| MAL | 0.1236 | 0.0004 | 0.0289 | 0.4897 | 0.0036 | 0.0098 |
| SATB2 | -0.0528 | 0.1319 | -0.0453 | 0.2191 | 0.0024 | 0.0066 |
| SIDT1 | 0.1259 | 0.0003 | 0.0139 | 0.7844 | 0.0017 | 0.0048 |
| HTT | -0.0251 | 0.4748 | -0.0503 | 0.4965 | 0.0013 | 0.0035 |
| IL4 | 0.0134 | 0.7020 | -0.0089 | 0.7981 | -0.0001 | -0.0003 |
| CD74 | -0.0251 | 0.4734 | 0.0640 | 0.2214 | -0.0016 | -0.0044 |
| HIST1H3H | 0.0667 | 0.0569 | 0.0922 | 0.0409 | 0.0061 | 0.0169 |
| CFB | -0.0251 | 0.4741 | 0.0168 | 0.6794 | -0.0004 | -0.0012 |
| AKAP12 | 0.0483 | 0.1688 | 0.1486 | 0.0009 | 0.0072 | 0.0197 |
| LPAL2 | -0.0715 | 0.0412 | 0.0267 | 0.4355 | -0.0019 | -0.0053 |
| PSMB1 | -0.0021 | 0.9515 | 0.0424 | 0.4669 | -0.0001 | -0.0002 |
| ITGB8 | 0.0602 | 0.0860 | -0.0539 | 0.2066 | -0.0032 | -0.0089 |
| MS4A2 | 0.0618 | 0.0780 | 0.0201 | 0.6733 | 0.0012 | 0.0034 |
| SCCPDH | -0.0010 | 0.9776 | 0.0360 | 0.4990 | -0.0000 | -0.0001 |
| C1orf198 | 0.1182 | 0.0007 | 0.0354 | 0.4390 | 0.0042 | 0.0115 |
| LMAN2L | -0.0815 | 0.0199 | -0.0249 | 0.4912 | 0.0020 | 0.0056 |
| DCUN1D4 | 0.0926 | 0.0082 | -0.2600 | 0.0001 | -0.0241 | -0.0663 |
| PDGFRA | 0.0365 | 0.2977 | 0.0429 | 0.3270 | 0.0016 | 0.0043 |
| MFAP3L | 0.0940 | 0.0073 | 0.0848 | 0.0992 | 0.0080 | 0.0219 |
| HIST1H4A | -0.0269 | 0.4425 | 0.0156 | 0.6490 | -0.0004 | -0.0012 |
| ABCA13 | -0.1958 | 0.0000 | 0.0657 | 0.2759 | -0.0129 | -0.0354 |
| FLJ43692 | 0.0438 | 0.2121 | 0.0145 | 0.6781 | 0.0006 | 0.0017 |
| SQLE | -0.0337 | 0.3366 | -0.1304 | 0.0006 | 0.0044 | 0.0121 |
| PPP2CB | 0.0489 | 0.1633 | 0.0641 | 0.1654 | 0.0031 | 0.0086 |
| CTTN | 0.1422 | 0.0000 | -0.0629 | 0.2616 | -0.0089 | -0.0246 |
| MMP8 | -0.1680 | 0.0000 | -0.0936 | 0.0920 | 0.0157 | 0.0433 |
| CD9 | -0.0339 | 0.3332 | -0.0206 | 0.6156 | 0.0007 | 0.0019 |
| GPR84 | -0.0980 | 0.0051 | -0.0703 | 0.0533 | 0.0069 | 0.0190 |
| FAM19A2 | -0.0978 | 0.0052 | 0.0328 | 0.3476 | -0.0032 | -0.0088 |
| SPRED1 | -0.0045 | 0.8971 | 0.0791 | 0.0333 | -0.0004 | -0.0010 |
| CETP | -0.0356 | 0.3100 | -0.0365 | 0.3285 | 0.0013 | 0.0036 |
| BPI | -0.2393 | 0.0000 | 0.0160 | 0.7623 | -0.0038 | -0.0105 |
| TGM2 | -0.0370 | 0.2911 | -0.0312 | 0.4298 | 0.0012 | 0.0032 |
| NGFRAP1 | 0.1193 | 0.0006 | -0.0289 | 0.5858 | -0.0034 | -0.0095 |
| Direct Effect | - | - | - | - | 0.3545 | 0.0000 |

**Table 3.** Product measure estimates in FHS dataset, subgroup 2 (BMI $\leq 25$, Age $> 50$). For Direct effect, the product is the estimation of indirect effect. Med Prop: Mediation proportion.

| Mediator | a | p.value(a) | b | p.value(b) | product | med prop |
|---|---|---|---|---|---|---|
| ZBTB40 | -0.0582 | 0.0064 | 0.0027 | 0.9439 | -0.0002 | -0.0004 |
| LAMC1 | 0.0078 | 0.7136 | -0.0521 | 0.0231 | -0.0004 | -0.0010 |
| DEDD | -0.0205 | 0.3364 | -0.0976 | 0.0002 | 0.0020 | 0.0050 |
| PTPN7 | -0.0918 | 0.0000 | 0.0255 | 0.3728 | -0.0023 | -0.0059 |
| MAL | 0.1919 | 0.0000 | -0.0428 | 0.0760 | -0.0082 | -0.0206 |
| SATB2 | -0.0649 | 0.0024 | -0.0006 | 0.9783 | 0.0000 | 0.0001 |
| SIDT1 | 0.0101 | 0.6357 | 0.0085 | 0.7716 | 0.0001 | 0.0002 |
| HTT | -0.0569 | 0.0077 | 0.0098 | 0.8086 | -0.0006 | -0.0014 |
| IL4 | 0.0466 | 0.0291 | 0.0000 | 0.9998 | 0.0000 | 0.0000 |
| CD74 | -0.1010 | 0.0000 | 0.1037 | 0.0012 | -0.0105 | -0.0263 |
| HIST1H3H | 0.1201 | 0.0000 | 0.0459 | 0.0613 | 0.0055 | 0.0138 |
| CFB | -0.0458 | 0.0320 | -0.0129 | 0.5602 | 0.0006 | 0.0015 |
| AKAP12 | 0.1179 | 0.0000 | 0.0887 | 0.0010 | 0.0105 | 0.0262 |
| LPAL2 | -0.0868 | 0.0000 | 0.0065 | 0.7396 | -0.0006 | -0.0014 |
| PSMB1 | -0.0608 | 0.0044 | -0.0594 | 0.0786 | 0.0036 | 0.0091 |
| ITGB8 | 0.1020 | 0.0000 | -0.0180 | 0.4437 | -0.0018 | -0.0046 |
| MS4A2 | 0.1893 | 0.0000 | 0.1208 | 0.0000 | 0.0229 | 0.0573 |
| SCCPDH | -0.0256 | 0.2314 | 0.0984 | 0.0024 | -0.0025 | -0.0063 |
| C1orf198 | 0.1856 | 0.0000 | 0.0761 | 0.0046 | 0.0141 | 0.0354 |
| LMAN2L | -0.0093 | 0.6631 | -0.0242 | 0.2431 | 0.0002 | 0.0006 |
| DCUN1D4 | 0.0599 | 0.0050 | -0.1219 | 0.0018 | -0.0073 | -0.0183 |
| PDGFRA | -0.0304 | 0.1544 | -0.0030 | 0.9065 | 0.0001 | 0.0002 |
| MFAP3L | 0.1203 | 0.0000 | 0.0550 | 0.0575 | 0.0066 | 0.0166 |
| HIST1H4A | -0.0240 | 0.2607 | -0.0155 | 0.4165 | 0.0004 | 0.0009 |
| ABCA13 | -0.2092 | 0.0000 | -0.0257 | 0.4920 | 0.0054 | 0.0135 |
| FLJ43692 | 0.0474 | 0.0263 | 0.0137 | 0.4916 | 0.0006 | 0.0016 |
| SQLE | -0.0626 | 0.0034 | -0.1392 | 0.0000 | 0.0087 | 0.0218 |
| PPP2CB | -0.0281 | 0.1885 | 0.0902 | 0.0007 | -0.0025 | -0.0063 |
| CTTN | 0.1416 | 0.0000 | -0.0679 | 0.0340 | -0.0096 | -0.0241 |
| MMP8 | -0.1514 | 0.0000 | -0.0853 | 0.0179 | 0.0129 | 0.0324 |
| CD9 | 0.0205 | 0.3363 | -0.0284 | 0.2219 | -0.0006 | -0.0015 |
| GPR84 | -0.0051 | 0.8128 | 0.0039 | 0.8551 | -0.0000 | -0.0000 |
| FAM19A2 | -0.0997 | 0.0000 | 0.0244 | 0.2384 | -0.0024 | -0.0061 |
| SPRED1 | -0.0610 | 0.0043 | 0.0466 | 0.0315 | -0.0028 | -0.0071 |
| CETP | 0.0440 | 0.0393 | -0.0333 | 0.1251 | -0.0015 | -0.0037 |
| BPI | -0.2516 | 0.0000 | 0.0474 | 0.1652 | -0.0119 | -0.0299 |
| TGM2 | 0.1257 | 0.0000 | -0.0534 | 0.0242 | -0.0067 | -0.0168 |
| NGFRAP1 | 0.1478 | 0.0000 | 0.0663 | 0.0187 | 0.0098 | 0.0246 |
| Direct Effect | - | - | - | - | 0.3673 | 0.0000 |

**Table 4.** Product measure estimates in FHS dataset, subgroup 3 (BMI $>$ 25, Age $\leq$ 62). For Direct effect, the product is the estimation of indirect effect. Med Prop: Mediation proportion.

| Mediator | a | p.value(a) | b | p.value(b) | Product | Med prop |
|---|---|---|---|---|---|---|
| ZBTB40 | 0.0767 | 0.0278 | -0.0334 | 0.5960 | -0.0026 | -0.0066 |
| LAMC1 | 0.1556 | 7.346e-06 | -0.0040 | 0.9156 | -0.0006 | -0.0016 |
| DEDD | 0.0106 | 0.7617 | -0.1440 | 0.0014 | -0.0015 | -0.0039 |
| PTPN7 | 0.0215 | 0.5381 | 0.0024 | 0.9581 | 0.0001 | 0.0001 |
| MAL | 0.1794 | 2.207e-07 | -0.0791 | 0.0581 | -0.0142 | -0.0367 |
| SATB2 | -0.0213 | 0.5409 | 0.0181 | 0.5957 | -0.0004 | -0.0010 |
| SIDT1 | 0.1410 | 4.950e-05 | 0.0281 | 0.5487 | 0.0040 | 0.0102 |
| HTT | 0.0437 | 0.2104 | 0.1135 | 0.0912 | 0.0050 | 0.0128 |
| IL4 | -0.0613 | 0.0786 | 0.0230 | 0.5107 | -0.0014 | -0.0036 |
| CD74 | 0.0468 | 0.1802 | 0.1011 | 0.0534 | 0.0047 | 0.0122 |
| HIST1H3H | 0.0723 | 0.0382 | 0.0845 | 0.0461 | 0.0061 | 0.0158 |
| CFB | -0.0341 | 0.3282 | -0.0275 | 0.4741 | 0.0009 | 0.0024 |
| AKAP12 | 0.0328 | 0.3472 | 0.1110 | 0.0108 | 0.0036 | 0.0094 |
| LPAL2 | 0.0714 | 0.0406 | -0.0801 | 0.0179 | -0.0057 | -0.0148 |
| PSMB1 | -0.0670 | 0.0547 | -0.0285 | 0.6184 | 0.0019 | 0.0049 |
| ITGB8 | -0.0781 | 0.0250 | -0.0214 | 0.6157 | 0.0017 | 0.0043 |
| MS4A2 | -0.0902 | 0.0097 | 0.0654 | 0.1880 | -0.0059 | -0.0152 |
| SCCPDH | -0.1214 | 0.0005 | 0.0810 | 0.1113 | -0.0098 | -0.0254 |
| C1orf198 | 0.1492 | 1.724e-05 | 0.0870 | 0.0657 | 0.0130 | 0.0335 |
| LMAN2L | 0.0007 | 0.9844 | -0.0275 | 0.4319 | -0.0000 | -0.0000 |
| DCUN1D4 | -0.0298 | 0.3929 | -0.1066 | 0.0678 | 0.0032 | 0.0082 |
| PDGFRA | -0.0122 | 0.7263 | -0.0115 | 0.7905 | 0.0001 | 0.0004 |
| MFAP3L | 0.0879 | 0.0116 | 0.0493 | 0.3119 | 0.0043 | 0.0112 |
| HIST1H4A | -0.0900 | 0.0098 | -0.0556 | 0.0982 | 0.0050 | 0.0129 |
| ABCA13 | -0.1555 | 7.412e-06 | -0.0290 | 0.6166 | 0.0045 | 0.0117 |
| FLJ43692 | -0.0179 | 0.6080 | -0.0199 | 0.5556 | 0.0004 | 0.0009 |
| SQLE | -0.0150 | 0.6673 | -0.0998 | 0.0083 | 0.0015 | 0.0039 |
| PPP2CB | -0.0052 | 0.8812 | 0.0677 | 0.1228 | -0.0004 | -0.0009 |
| CTTN | 0.1992 | 8.284e-09 | -0.1207 | 0.0247 | -0.0240 | -0.0621 |
| MMP8 | -0.1106 | 0.0015 | -0.0238 | 0.6788 | 0.0026 | 0.0068 |
| CD9 | -0.0517 | 0.1381 | -0.0212 | 0.5897 | 0.0011 | 0.0028 |
| GPR84 | -0.0708 | 0.0424 | -0.0345 | 0.3317 | 0.0024 | 0.0063 |
| FAM19A2 | -0.1226 | 0.0004 | 0.0513 | 0.1364 | -0.0063 | -0.0162 |
| SPRED1 | -0.0485 | 0.1645 | 0.0407 | 0.2699 | -0.0020 | -0.0051 |
| CETP | 0.1013 | 0.0036 | 0.0225 | 0.5510 | 0.0023 | 0.0059 |
| BPI | -0.1916 | 3.037e-08 | -0.0325 | 0.5439 | 0.0062 | 0.0161 |
| TGM2 | 0.0449 | 0.1977 | 0.0005 | 0.9887 | 0.0000 | 0.0001 |
| NGFRAP1 | 0.0956 | 0.0060 | 0.0611 | 0.1935 | 0.0058 | 0.0151 |
| Direct Effect | - | - | - | - | 0.3814 | 0.0000 |

**Table 5.** Product measure estimates in FHS dataset, subgroup 4 (BMI $>$ 25, Age $>$ 62). For Direct effect, the product is the estimation of indirect effect. Med Prop: Mediation proportion.

Xingyu Li, Qing Liu, Xun Jiang, Hong Amy Xia, Brian P Hobbs, and Peng Wei. Modeling heterogeneous mediation effects in survival analysis via an interpretable m-learner framework. *arXiv preprint arXiv:2603.13464*, 2026.

Laurens van der Maaten and Geoffrey Hinton. Visualizing data using t-sne. *Journal of machine learning research*, 9(Nov):2579–2605, 2008.
